# Synaptotagmin 13 orchestrates pancreatic endocrine cell egression and islet morphogenesis

**DOI:** 10.1101/2021.08.30.458251

**Authors:** Mostafa Bakhti, Aimée Bastidas-Ponce, Sophie Tritschler, Marta Tarquis-Medina, Eva Nedvedova, Katharina Scheibner, Jessica Jaki, Perla Cota, Ciro Salinno, Karsten Boldt, Stefanie J. Willmann, Nicola Horn, Marius Ueffing, Ingo Burtscher, Fabian J. Theis, Ünal Coskun, Heiko Lickert

**Affiliations:** Institute of Diabetes and Regeneration Research, Helmholtz Zentrum München, 85764 Neuherberg, Germany; German Center for Diabetes Research (DZD), D-85764 Neuherberg, Germany; Institute of Computational Biology, Helmholtz Zentrum München, D-85764 Neuherberg, Germany; Technical University of Munich, School of Life Sciences Weihenstephan, 85354 Freising, Germany; Technische Universität München, School of Medicine, 81675 München, Germany; Paul Langerhans Institute Dresden of the Helmholtz Zentrum Munich at the University Clinic Carl Gustav Carus, TU Dresden, Dresden, Germany; Institute for Clinical Chemistry and Laboratory Medicine, University Hospital Carl Gustav Carus of Technische Universität Dresden, Fetscher Str. 74, 01307 Dresden; Institute for Ophthalmic Research, Center for Ophthalmology, University of Tübingen, 72076 Tübingen, Germany; Technical University of Munich, Department of Mathematics, 85748 Garching b. Munich, Germany; SOTIO a.s, Jankovcova 1518/2, 17000 Prague, Czech Republic

**Author notes:** Correspondence (M.B.), (H.L.). These authors contributed equally (Mostafa Bakhti, Aimée Bastidas-Ponce).

**Keywords:** Synaptotagmin 13, endocrine precursors, morphogenesis, egression, migration, polarity, cytoskeleton, acetylated tubulin, endocytosis, basement membrane

## Abstract

Epithelial cell egression is important for organ development, but also drives cancer metastasis. Better understandings of pancreatic epithelial morphogenetic programs generating islets of Langerhans aid to diabetes therapy. Here we identify the Ca^2+^-independent atypical Synaptotagmin 13 (Syt13) as a key driver of endocrine cell egression and islet formation. We detected upregulation of Syt13 in endocrine precursors that correlates with increased expression of unique cytoskeletal components. High-resolution imaging reveals a previously unidentified apical-basal to front-rear repolarization during endocrine cell egression. Strikingly, Syt13 interacts with acetylated tubulin and phosphatidylinositol phospholipids and localizes to the leading-edge of egressing cells. Knockout of Syt13 impairs endocrine cell egression and skews the α- to-β-cell ratio. Mechanistically, Syt13 regulates endocytosis to remodel the basement membrane and cell-matrix adhesion at the leading-edge of egressing endocrine cells. Altogether, these findings implicate an unexpected role of Syt13 in regulating cell polarity to orchestrate endocrine cell egression and islet morphogenesis.

## Introduction

The pancreatic endocrine cells (α-, β-, δ-, Ɛ-, and PP-cells) regulate glucose homeostasis. How endocrine lineage segregation in interlinked with morphogenesis is not well understood. During endocrinogenesis, a subset of multi-/bipotent progenitors, that reside in the epithelium, gradually express the transcription factor (TF) Neurogenin 3 (Neurog3; hereafter called Ngn3) to sequentially generate endocrine progenitors (Ngn3^low^) and precursors (Ngn3^high^) ^1^. The Ngn3^high^ precursors give rise to different endocrine cell types via a transient intermediate expressing the transcription factor (TF) Fev ^2^ (Figure 1A). The stepwise lineage formation is tightly connected with tissue morphogenesis, in which endocrine cells egress from the ductal epithelium (also referred to endocrine cell delamination) and cluster into the proto-islets ^3–5^. These morphogenetic events are initiated after Ngn3 induction and followed by the reduction of the apical plasma membrane (PM) area to form a tether structure, which ultimately undergoes abscission ^6–8^. Cytoskeletal dynamics and small GTPases such as Cdc42 and Rac1 are involved in these stepwise events and coordinate local endocrine cell egression to form proto-islets described as the peninsular model ^9–11^. However, whether asymmetric cell division ^4,12^, endocrine cell migration ^13,14^ and epithelial-to-mesenchymal transition (EMT) ^8,15^ mediate endocrine cell egression from the pancreatic epithelium is still a matter of debate. Moreover, the upstream morphogenetic drivers orchestrateing cell and cytoskeletal dynamics during endocrine cell egression have not been identified. Further, the molecular mechanisms enforcing egressing cells to push or break through the basal lamina are unknown. Deciphering these mechanisms aids to tissue engineer islets for cell-replacement therapy but also to identify novel targets to intervene on epithelial cell dissemination during cancer metastasis.

**Figure 1.**
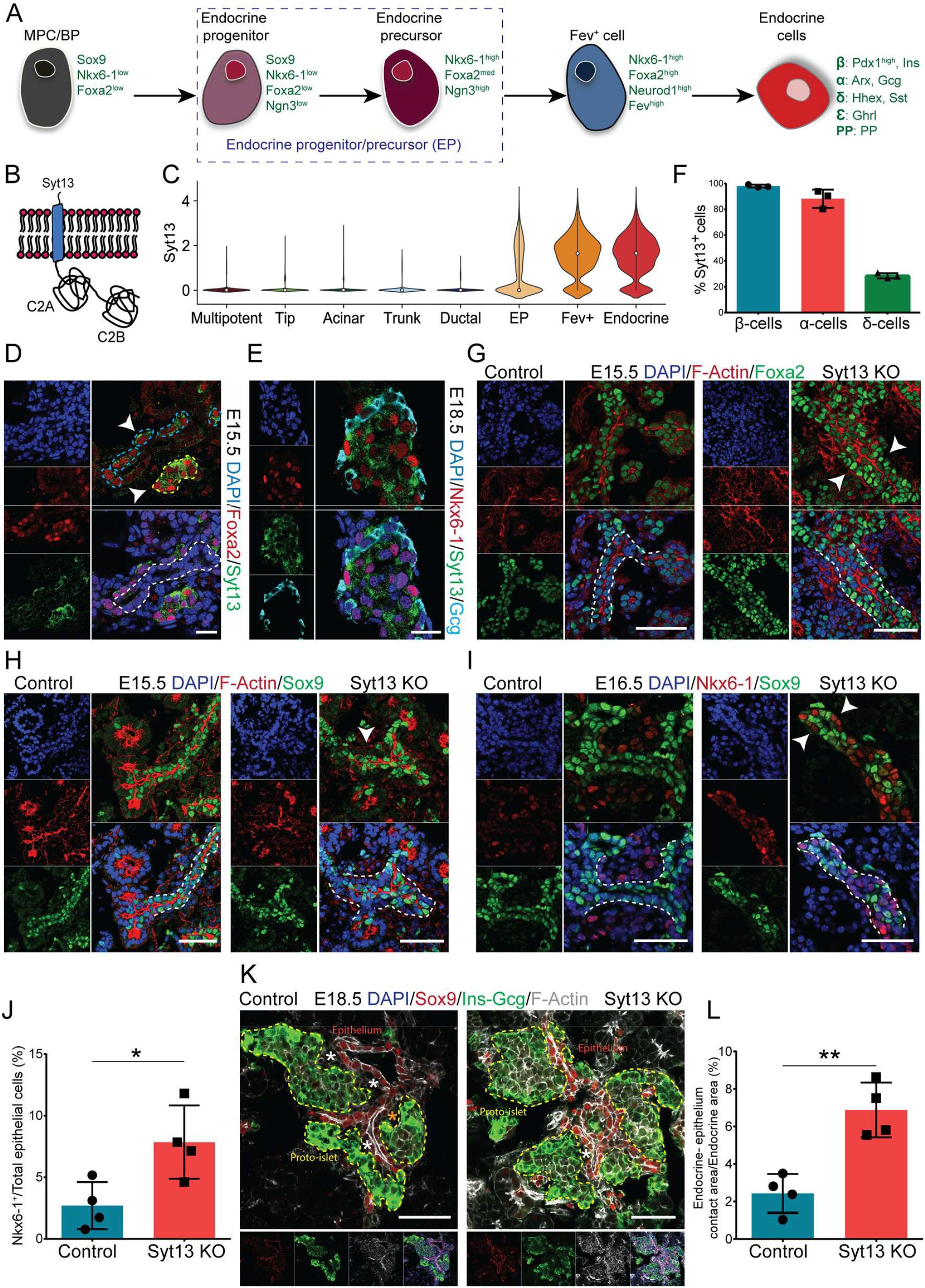
Loss of Syt13 hampers endocrine cell egression. (A) Schematic picture of stepwise endocrine lineage formation. MPC, multipotent progenitor cells; BP, bipotent progenitors. (B) Schematic representation of Syt13 protein. (C) Violin plot showing Syt13 expression in pancreatic lineages in scRNA-seq data of mouse endocrinogenesis (E12.5 – E15.5). (D) Syt13 expression in pancreatic epithelial cells (Foxa2^high^, arrowhead). (E) Expression of Syt13 in β-cells (Nkx6-1^high^) and α-cells (Glucagon^+^). (F) The percentage of β-, α- and δ-cell expressing Syt13 in pancreatic sections at E18.5. (G) A multi-layer epithelium (arrowheads) appears in Syt13KO pancreata. The extra layer is composed of Foxa2^high^ cells. (H, I) The cells in the extra epithelial layer in Syt13 KO pancreata are negative for Sox9 (arrowhead in H) but express high levels of Nkx6-1 (arrowheads in I). (J) Quantification of the number of Nkx6-1^high^/Sox9^-^ cells within or associated with the epithelium. (K) IHC analysis of the proto-islet position to the epithelium. * shows the area where proto-islets are not in direct contact with the nearby epithelium. (L) Quantification of contact area between proto-islets and the epithelium in Syt13 KO and control mice at E18.5. Scale bar 20 µm (D); 50 µm (E, G-I, K). (*P < 0.05; **P < 0.01; t-test). Data are represented as mean ± SD. See also Figure S1 and S2.

Here, we report Synaptotagmin 13 (Syt13) (Figure 1B) as a major regulator of endocrine cell egression and islet morphogenesis during endocrinogenesis. Syt13 is an atypical member of the Synaptotagmin (SYT) family of membrane trafficking proteins, which are known to be involved in intracellular vesicle trafficking and exocytosis. This protein family consists of 17 members and each possesses a transmembrane sequence connected to two lipid binding cytoplasmic domains (C2A and C2B), responsible for docking and fusion of the carrier vesicles to the target membrane ^16,17^. Compared to well-known members, such as Syt1 and Syt2, Syt13 does not have an N-terminal sequence preceding the transmembrane region, and does not interact with membranes in a calcium (Ca^2+^)-dependent manner ^18,19^. Further, although classical Syt proteins are involved in vesicle exocytosis, the cellular and molecular function of atypical members, such as Syt13, is less described.

By combining mouse genetics, high-resolution cell biology and single cell mRNA profiling, we identified a critical function of Syt13 in pancreatic endocrine morphogenesis. Our data identified apical-basal to front rear polarization during endocrine cell egression, which was accompanied by upregulation of molecular components involved in cell motility and cytoskeletal dynamics. Importantly, we uncovered Syt13 as a leading-edge protein that remodels the basement membrane and cell-matrix adhesion during endocrine morphogenesis. Together, our findings provide a detailed mechanistic description of how endocrine precursors leave the pancreatic epithelium and discovered Syt13 as a polarity protein and morphogenetic regulator during islet formation.

## Results

### Syt13 function is required for endocrine cell egression and migration

In a global gene expression profiling during pancreas development, we previously identified Syt13 to be expressed during the peak of endocrinogenesis ^20^. To further pinpoint the temporal and cell-type specific expression pattern of *Syt13* mRNA, we leveraged single-cell RNA-sequencing (scRNA-seq) data from mouse embryonic pancreatic epithelial cells that were sampled from E12.5 to E.15.5 ^21^. *Syt13* expression was specific to endocrine progenitors/precursors (EPs) and lineage cells (Figure 1C), and together with *Syt7* first expressed of all *SYT* family members during endocrinogenesis (Figure S1A). Likewise, we found *SYT13* and *SYT7* to be early-onset *SYT* genes in human *in vitro* stem cell differentiation toward endocrine lineage ^22^ (Figure S1B). Next, we confirmed Syt13 protein synthesis and localization in cells expressing high levels of TF Foxa2 (Foxa2^high^) leaving the ductal epithelium (Figure 1D, blue dashed lines) or clustered in proto-islets (Figure 1D, yellow dashed line). Further analysis indicated that Syt13 protein expression was restricted to the endocrine lineage and it was synthesized in a major fraction of embryonic α- and β-cells, but only in a minor fraction of δ-cells (Figure 1E and 1F). To uncover the cellular and molecular function of Syt13, we generated a Sy13 knockout (KO) mouse line using the genetrap targeted mESCS (EUCOMM), in which the critical exon 2 was removed. This strategy resulted in the generation of a Flox (F)-deleted (FD) allele and whole-body *Syt13*^*FD/FD*^ (Syt13 KO) mice (Figure S1C-F). Mendelian ratio of heterozygous (*Syt13*^*+/FD*^) intercrosses indicated no lethality of *Syt13*^*FD/FD*^ at embryonic but at prenatal stages (Figure S1G). As the heterozygous mice showed similar phenotype to the wild type (WT) animals, we refer to both genotypes hereafter as controls. We detected similar size and weight of control and *Syt13*^*FD/FD*^ embryos at E18.5 (Figure S1H and S1I). Gross morphology of control and *Syt13*^*FD/FD*^ pancreata were comparable at E18.5 (Figure S1J). Moreover, we found no evident histological alterations in the *Syt13*^*FD/FD*^ pancreatic epithelium at E12.5 and E13.5 (Figure S2A and S2B). Further analysis revealed no changes in apical-basal polarity in *Syt13*^*FD/FD*^ ductal epithelial and acinar cells (Figure S2C and S2D). These data reveal normal ductal and acinar differentiation and morphogenesis upon *Syt13* deletion and indicate that Syt13 expression and function is highly specific to the endocrine compartment.

Starting from E14.5, striking alterations in epithelial organization appeared in the Syt13 KO pancreata. While the pancreatic epithelium of control mice consisted of a single-layer of Foxa2^low^ cells, the Syt13 KO pancreata contained a multi-layer epithelium. The extra layers were mainly Foxa2^high^ cells (Figure 1G and S2E) and negative for the bipotent/ductal cell marker, Sox9 (Figure 1H), suggesting a defect during endocrine lineage acquisition. In support of this, we found an increased number of retained EPs/endocrine cells (Nkx6-1^high^/Sox9^-^) within the *Syt13*^*FD/FD*^ compared to the control epithelium (Figure 1I and 1J). Moreover, we detected an increased direct attachment between the proto-islets and the epithelium in *Syt13*^*FD/FD*^ compared to control pancreata (Figure 1K and 1L). Additionally, the typical proto-islet rearrangement, in which α-cells are at the periphery and β-cells are at the core, was disrupted in Syt13 KO pancreata (Figure S2F). To dissect at which cell state Syt13 regulates endocrine cell egression, we generated two lineage and cell type-specific Syt13-conditional KO (CKO) mice using *Ngn3*^*Cre/+*^ and *Ins1*^*Cre/+*^ Cre-driver lines to delete Syt13 specifically in EPs and the β-cell lineage, respectively. We found an impairment in β-cell egression/migration in *Ngn3* ^*Cre/+*^; *Syt13*^*F/FD*^, but not in *Ins1* ^*Cre/+*^; *Syt13*^*F/FD*^ animals (Figure S2G and S2H). We conclude that Syt13 function is crucial for endocrine cell egression in EPs, but not in newly formed β-cells.

### High expression levels of *Syt13* in endocrine precursors and *Fev*^+^ cells prime β-cell fate

We next dissected the expression dynamics of Syt13 in endocrine lineages using the scRNA-seq data ^21^. Syt13 expression levels increased from *Ngn3*^low^ progenitors to *Ngn3*^high^ precursors (Figure 2A). Moreover, *Ngn3*^high^ precursors from E14.5 and E15.5 had higher levels of *Syt13* compared to those from E12.5 and E13.5 (Figure 2B). To investigate the possible link between Syt13 expression levels and different endocrine cell fates, we divided *Ngn3*^high^ precursors from E12.5-15.5 into two clusters expressing high levels (*Syt13*^high^) and no or low levels of *Syt13* (*Syt13*^low/-^) (Figure 2C). The fraction of *Syt13*^high^ cells increased from E12.5 to E15.5 (Figure 2D). Differential gene expression analysis between *Syt13*^high^ and *Syt13*^low/-^ EPs identified several previously reported EP-signature genes ^21^ (Figure 2E and Table S2). The majority of signature genes, which likely associate with the β-cell fate and include *Neurod2, Sult2b1* and *Upk3bl*, were highly expressed in the *Syt13*^high^ precursors, whereas genes, which are possibly linked to the α-cell fate including *Dll1, Rsad2* and *Rasgrp3*, were highly expressed in *Syt13*^low/-^ precursors (Figure 2F). Furthermore, the expression levels of *Pax4, Gck, Nkx6-1* and *Nkx2-2* were higher in *Syt13*^high^ than *Syt13*^low/-^ precursors, suggesting that *Syt13*^high^ *Ngn3*^high^ cells are primed for β-cell fate allocation (Figure 2G).

**Figure 2.**
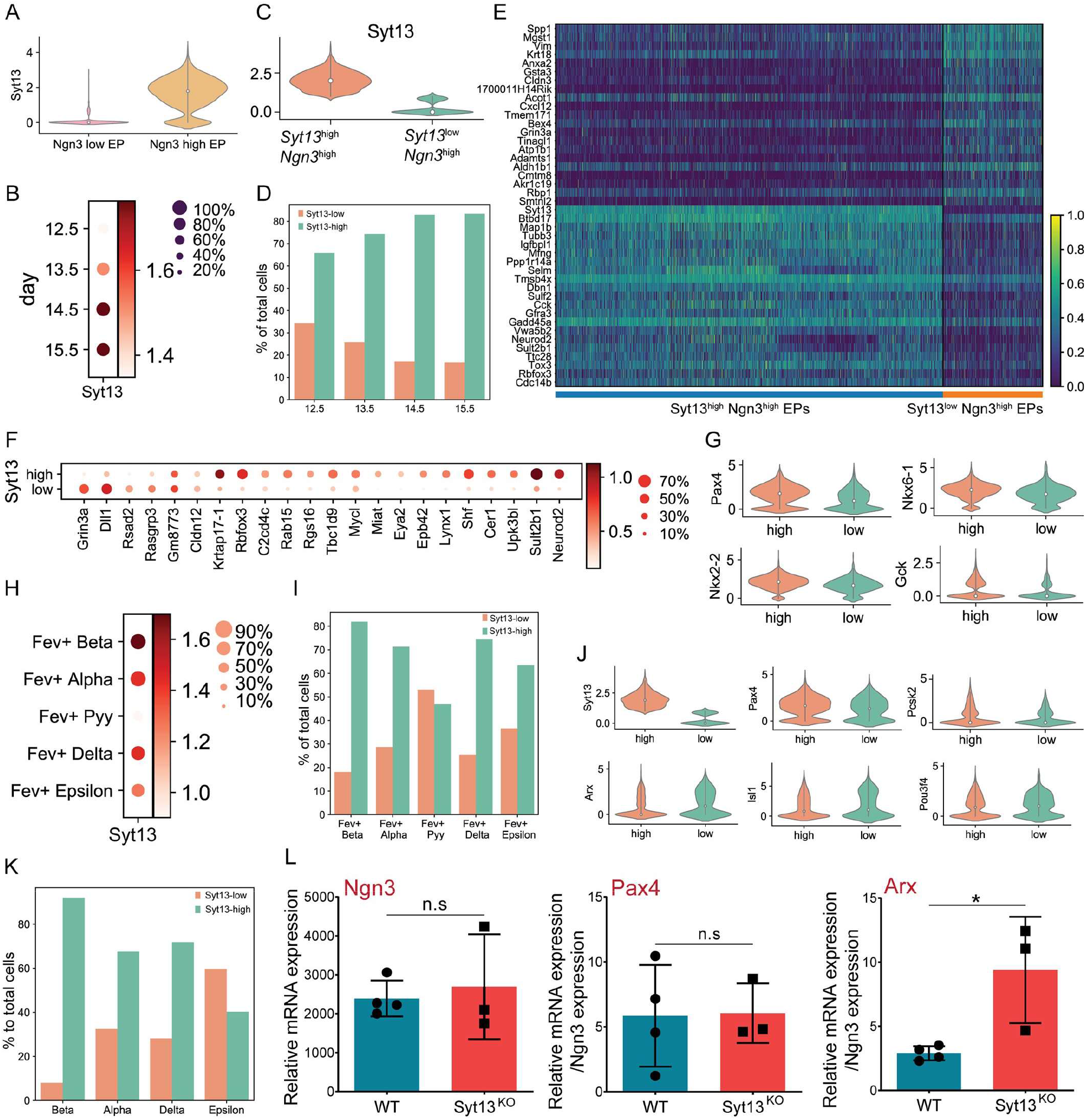
Increased expression levels of *Syt13* associate with the β-cell program. (A) Violin plot of *Syt13* expression levels in endocrine progenitors and precursors. (B) Dotplot of *Syt13* expression in endocrine precursors at different developmental stages. (C) Violin plot shows the dividing of the endocrine precursors into *Syt13*^high^ and *Syt13*^low/-^ clusters. (D) The percentage of *Syt13*^high^ *Ngn3*^high^ and *Syt13*^low/-^ *Ngn3*^high^ cells at different developmental stages. (E) Headmap of top 21 differentially expressed genes in *Syt13*^high^ and *Syt13*^low/-^ precursors. (F) Dotplot indicating the differential expression of EP-signature genes in *Syt13*^high^ and *Syt13*^low/-^ precursors. (G) Violin plots showing the expression of β-cell specific genes in *Syt13*^high^ and *Syt13*^low/-^ precursors. (H) Dotplot of *Syt13* expression in different *Fev*^+^ cells from E12.5-E15.5. (I) The percentage of *Syt13*^high^ and *Syt13*^low/-^ cells in different *Fev*^+^ cell types. (J) Violin plots showing the expression of β- and α-cell specific genes in *Syt13*^high^ and *Syt13*^low/-^ *Fev*^+^ cells. (K) The percentage of *Syt13*^high^ and *Syt13*^low/-^ cells in different hormone^+^ endocrine cell types from E12.5-E15.5. (L) qPCR analysis of NVF^+^ cells sorted from pancreatic epithelium from NVF;control and NVF;Syt13 KO mice at E15.5. (n.s, non-significant; *P < 0.05; t-test). Data are represented as mean ± SD. See also Figure S3.

We then analyzed Syt13 expression in *Fev*^+^ cells and found higher expression levels in *Fev*^+^β compared to other *Fev*^+^ endocrine subtypes (Figure 2H). Additionally, the percentage of *Syt13*^high^ cells was higher in *Fev*^+^β cells than in *Fev*^+^α, *Fev*^+^δ and *Fev*^+^ε and *Fev*^+^Pyy cells. (Figure 2I). Next, we splitted all *Fev*^+^ cells into *Syt13*^high^ and *Syt13*^low/-^ cells and performed differential gene expression analysis (Figure S3A and Table S2). *Pax4* and *Pcsk2* were upregulated in *Syt13*^high^, while *Arx, Isl1* and *Pou3f4* were upregulated *Syt13*^low/-^ *Fev*^+^ cells (Figure 2J). These data indicate that higher expression levels of *Syt13* in *Fev*^+^ cells associate with β-cell program. Consistent with *Syt13* expression in EPs and *Fev*^+^ cells, the fraction of *Syt13*^high^ cells was highest in β-cells of all different hormone^+^ endocrine subtypes (Figure 2K). Moreover, partition-based graph abstraction (PAGA) analysis showed high connectivity between *Syt13*^high^ precursors/*Fev*^+^ cells, which indicates that *Syt13*^high^ precursors/*Fev*^+^ cells transition to β-cells (Figure S3B). In line with these findings, in scRNA-seq data of human fetal pancreas ^23^ we found higher *SYT13* expression in fetal β-cells and their precursors compared to fetal α-cells and their precursors (Figure S3C). To support the findings from scRNA-seq analysis, we performed qPCR analysis of FACS isolated Ngn3^+^ cells from E15.5 *NVF*; *Syt13*^*FD/FD*^ pancreata, which were obtained through crossing *Syt13*^*+/FD*^ mice with homozygous Ngn3-Venus fusion (NVF) reporter mouse line. We found comparable levels of *Ngn3* in the control and *Syt13*^*FD/FD*^ Ngn3^+^ cells. Furthermore, an increased expression level of *Arx* but not *Pax4* was observed in the *Syt13*^*FD/FD*^ Ngn3^+^ cells compared to the controls (Figure 2L). Collectively, these analyses demonstrate that high physiological levels of *Syt13* mRNA expression in endocrine precursors and *Fev*^+^ cells correlates with β-cell fate acquisition.

### Lack of Syt13 is dispensable for endocrine induction but reduces β-cell specification

To validate the results from the scRNA-seq data, we examined the impact of Syt13 knockout on endocrine lineage formation. We found comparable numbers of Ngn3^+^ cells from control and Syt13 KO mice at E13.5-15.5 (Figure 3A and 3B). FACS analysis also revealed similar percentage of Ngn3^+^ cells in *NVF*; *Syt13*^*FD/FD*^ and control pancreata at E15.5 (Figure 3C). In addition, Syt13 KO and control pancreata contained a comparable fraction of cells expressing the pan-endocrine cell marker, Chromogranin A (ChgA) (Figure 3D and 3E). Next, we quantified the number of α- and β-cells at E14.5-16.5 and E18.5. A significant increase in the number of α-cells at the expense of β-cells in Syt13 KO pancreata was detected (Figure 3F and 3G). Importantly, we found that the deletion of Syt13 in EPs but not in the newly generated insulin-expressing cells resulted in an increased α- to β-cell ratio (Figure 3H-J and S3D). Collectively, these results support the findings from the *in vivo* scRNA-seq data and further indicate that Syt13 acts downstream of Ngn3 and its function in EPs is essential for β-cell fate allocation. Therefore, Syt13 regulates endocrine cell egression and allocation at EP/Fev^+^ state but not after β-cell specification.

**Figure 3.**
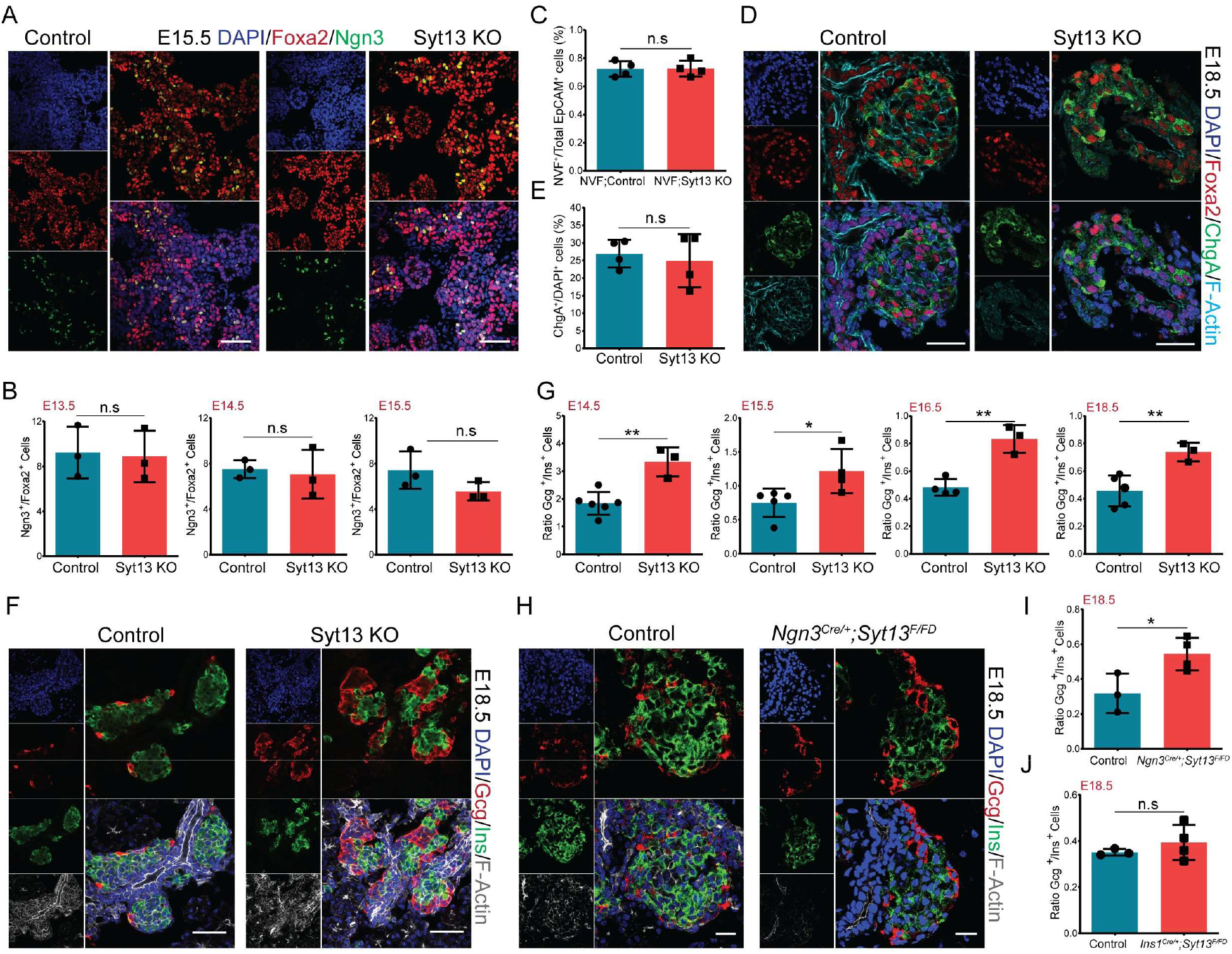
Loss of Syt13 reduces β-cell specification. (A) IHC analysis showing Ngn3^+^ cells in control and Syt13 KO mice. (B) Quantification of the percentage of Ngn3^+^ cells. (C) Percentage of NVF (Ngn3^+^) cells in the sorted epithelial cells from E15.5 pancreata. (D) IHC analysis of ChgA in pancreatic sections. (E) Quantification of percentage of ChgA^+^ cells in E18.5 pancreata. (F) IHC analysis of pancreatic sections from control and Syt13 KO mice. (G) Quantification of the α-/β-cells ratio in pancreatic sections from control and Syt13 KO mice. (H) IHC analysis of pancreatic sections from control and *Ngn3*^*Cre/+*^*;Syt13*^*F/FD*^ mice. (I, J) Quantification of the α-/β-cells ratio in *Ngn3*^*Cre/+*^*;Syt13*^*F/FD*^ and *Ins1*^*Cre/+*^*;Syt13*^*F/FD*^ pancreata, respectively. Scale bar 50 µm (A, D, F); 20 µm (H). (n.s, non-significant; *P < 0.05; **P < 0.01; t-test). Data are represented as mean ± SD. See also Figure S3.

### Syt13 localizes at the leading-edge of egressing endocrine cells

To decipher the cellular processes by which Syt13 coordinates endocrine cell egression and islet morphogenesis, we first explored the intracellular localization of the protein. The onset of *Syt13* mRNA expression occurs in the epithelium-residing EPs (Figure 1C). Therefore, we first assessed the localization of this protein in Syt13-overexpressing Madin-Darby Canine Kidney (MDCK) cells cultured in a 3D condition as a model for an apical-basal polarized epithelium. Remarkably, Syt13 was specifically localized at the apical domain of polarized epithelial cysts (Figure 4A). Proximity-dependent biotin identification (BioID) further identified the close proximity of Syt13 with several apical polarity determinants including aPKC, Ezrin, EBP50 and Merlin (Figure 4B). Surface biotinylation assay indicated the incorporation of Syt13 within the PM (Figure 4C). Further, structure-function study unveiled the requirement of both cytoplasmic C2A and C2B domains for integration and apical PM localization of the Syt13 protein (Figure 4D, E and S4A). This PM localization was different to Syt1 and Syt2, which are mainly enriched in the synaptic vesicles and mediate exocytosis. This suggests a distinct function of Syt13 compared to typical Syt proteins.

**Figure 4.**
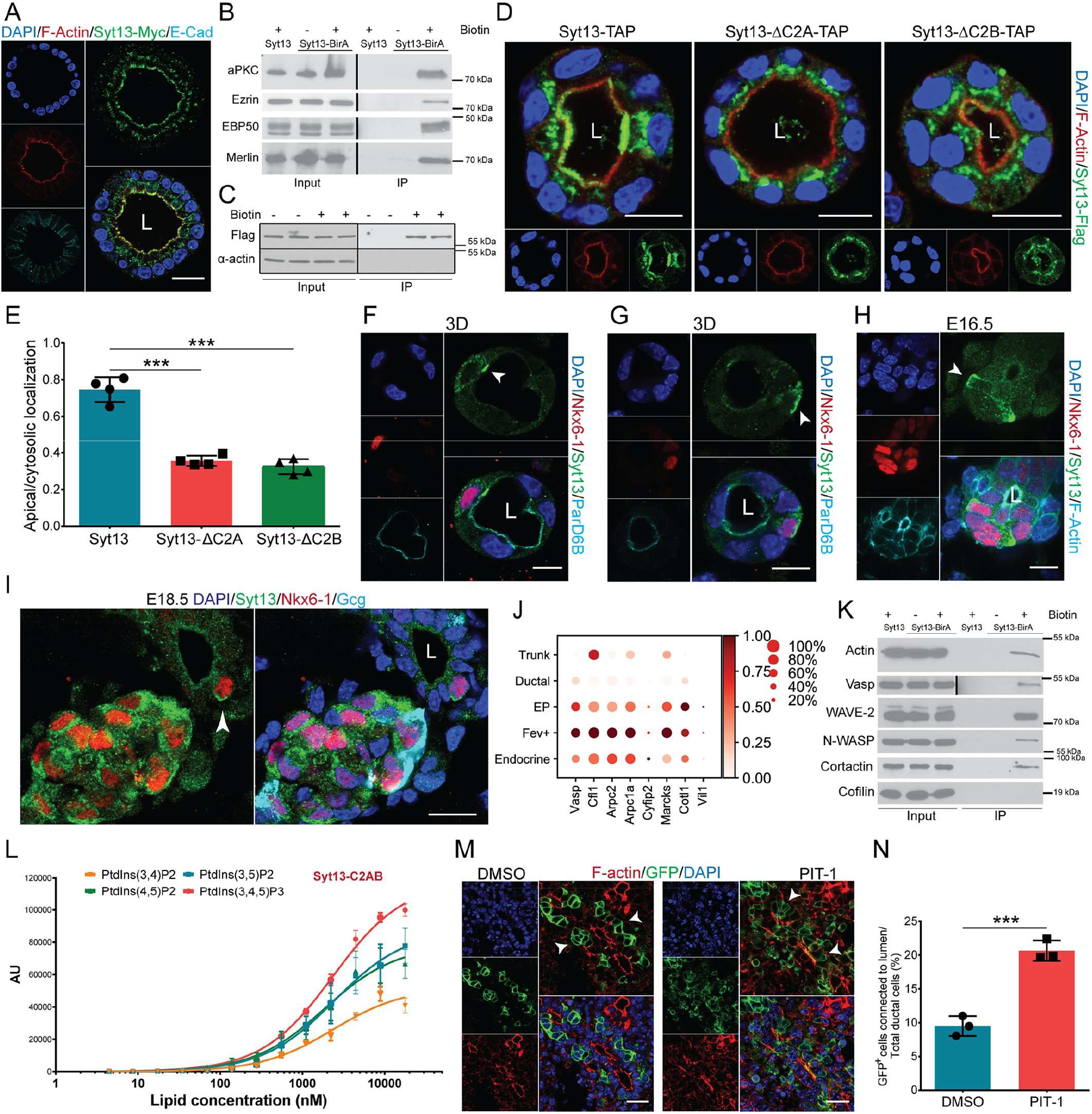
Syt13 localizes at the apical PM of epithelial and leading-edge of egressing endocrine cells. (A) Syt13 localizes at the apical PM in polarized MDCK cells. (B) Western blot showing the close proximity of Syt13 with different apical proteins in MDCK cells. (C) Surface biotinylation assay showing Syt13 localization to the plasma membrane. (D) Apical localization of Syt13 full-length and the C2 truncated variants. (E) Quantification of apical localization of Syt13 protein variants. (F) Syt13 localizes at the apical PM (arrowhead) in differentiated endocrine cells fully residing within the epithelium. (G, H) Syt13 localizes at the leading-edge (arrowhead) of egressing endocrine cells in 3D epithelial cysts (G) and *in vivo* (H). (I) Polarized Syt13 localization (arrowhead) in the single migrating endocrine cells. (J) Dot plot discloses the expression levels of the lamellipodium genes at different stages of mouse endocrinogenesis. (K) Western blot revealing close proximity of Syt13 protein with actin-based membrane protrusion proteins in MDCK cells. (L) Binding of purified Syt13 protein to 100 nm sized LUVs containing different species of phosphatidylinositol phospholipids (POPC/cholesterol/phosphoinositide 65/30/5 mol %) revealed by electrochemiluminescence-based immunoassay (EIA). (M, N) Treatment of explant culture of E13.5 *Ngn3*^*Cre/+*^; *ROSA26*^*mTmG/mTmG*^ pancreata with PIP3 inhibitor PIT-1 for 48 h, impairs endocrine cell egression. Arrowheads indicate endocrine cells. L, lumen. Scale bar 10 µm (D, F-H); 20 µm (A, M); 50 µm (I). (***P < 0.001; t-test). Data are represented as mean ± SD. See also Figure S4.

Next, we explored the cellular localization of Syt13 during endocrinogenesis. In newly generated endocrine cells, which were still fully integrated within the pancreatic epithelium, Syt13 was detected at the apical PM domain (Figure 4F and S4B). Importantly, during endocrine cell egression, Syt13 was mainly accumulated at the basal side of the ductal epithelium, which becomes the front of the precursors that leave the epithelium (Figure 4G and 4H). This localization pattern was also identified in cells, which appear to migrate out of the epithelium towards the proto-islet clusters (Figure 4I). Along this line, we found increased expression levels of several lamellipodium-related genes including *Vasp, Cotl1, Vil1, Arpc2, Cyfip2* and *Marcks* in *Syt13*^high^ compared to *Syt13*^low/-^ precursors (Figure S4C). Expression of these genes started in EPs and peaked in *Fev*^+^ cells (Figure 4J), which stresses that these cell states are highly motile. Additionally, we identified close proximity of Syt13 with several proteins including Actin, Vasp, Cortactin, the Arp2/3 regulators WAVE2 and N-WASP, but not with Cofilin (Figure 4K), further indicating leading-edge localization of Syt13.

One hallmark of directed cell migration is increased phosphatidylinositol triphosphate (PtdIns(3,4,5)P3 or PIP3) levels at the cell leading-edge. To investigate whether Syt13 interacts with phosphatidylinositol phospholipids, we performed an *in vitro* lipid-binding analysis of purified recombinant Syt13 protein variants (Figure S4D) to large unilamellar vesicles (LUVs). While C2A domain showed low binding to LUVs, C2B and C2AB domains exhibited binding preference towards PIP3, PtdIns(3,5)P2, PtdIns(4,5)P2 and with lower degree with PtdIns(3,4)P2 (Figure 4L and S4E-H). These data indicate that Syt13 directly interacts with PIP2 and PIP3 phosphoinositides mainly through its C2B cytoplasmic domain. Next, we tested if inhibition of PIP3 formation or function mimics the Syt13 action during endocrine cell egression. To test this, we used explant cultures from the lineage tracing *Ngn3*^*Cre/+*^; *ROSA26*^*mTmG/mTmG*^ pancreata and treated them with the PIP3/protein binding inhibitor (PIT-1) or the PI3K inhibitor (LY294002). Administration of both inhibitors resulted in striking impairments in the endocrine cell egression (Figure 4M, N and S4I). Overall, these data demonstrate that Syt13 localizes at the leading-edge of egressing cells to regulate endocrine cell repolarization along the front-rear axis (Figure 1).

### Syt13 levels correlate with the expression of unique molecular components of cell motility and cytoskeleton in endocrine precursors

To identify the molecular components linked with Syt13 functions during endocrine cell egression, we first analyzed the expression dynamics of EMT-related genes. We found low and transient expression of *Snail1/2* in EPs and a transient downregulation of *E-cadherin (Cdh1)* in *Fev*^+^ cells that did not correlate with increased expression of *N-cadherin (Cdh2)* at this cell state. Together with the absence of other classical EMT marker genes (Figure S5A), this suggests that endocrine cell egression is not mediated by a complete EMT program ^24^. Pathway enrichment analysis of differentially expressed genes between *Syt13*^high^ and *Syt13*^low/-^ precursors showed that increased *Syt13* expression in precursors coincided with upregulation of cell migration, actin and microtubule (MT) cytoskeletal organization, membrane protrusions and intracellular transport pathways (Figure 5A and Table S2). In *Syt13*^high^ precursors several genes that have been previously shown to regulate cell migration in different cellular systems and include *Tgfbr1, Cd40, Fgf18, Ret, Fer, Gfra3, Arhgef7, Slit1, Dok4, Olfm1* and *Lingo1* were highly expressed (Figure 5B and Table S2). Notably, the expression of most of these genes peaked at EPs or *Fev*^*+*^ cells, which indicates increased cell motility at these cell states (Figure S5B). Also several genes involved in actin and MT cytoskeletal rearrangement were differentially expressed. Tubulins (*Tuba1a, Tuba4a, Tubb3*), MT organizers (*Mtcl1, Mapre3, Kif26a*), MT-associated trafficking proteins (*Map1a, Map1b, Dynll2, Dync1i1, Kif5b*), actin remodeling proteins (*Tmsb4x, Dbn1, Scin, Vil1, Mical2*) and the cytoskeletal linker protein *dystonin (Dst)* were increased in *Syt13*^high^ compared to *Syt13*^low^ precursors (Figure 5C and Table S2). Remarkably, the expression onset of most of these genes was at EP state, which highlights their possible unique function in endocrine repolarization and dynamics (Figure S5B). Among these, we confirmed increased protein expression of α-tubulin (Tuba1a, Tuba4a), β3-tubulin (Tubb3), MT-associated protein 1b (Map1b), dystonin and Debrin-1 (Dbn1) during endocrinogenesis (Figure 5D and S5C). Moreover, using a newly established 2D (Figure S5D) and 3D culture systems, we disclosed co-expression of Syt13 protein with α-tubulin, β3-tubulin, Map1b, dystonin and Debrin-1 (Figure 5E and S5E). Interestingly, scRNA-seq data of human *in vitro* stem cell differentiation to EPs and the endocrine lineage validated upregulation of *MAP1B, DST* and *DBN1* in human endocrinogenesis ^22^. Moreover, these genes were increased in stem-cell derived β-cells (SC-β) compared to SC-α cells (Figure 5F and S5F). Finally, MAP1B, β3-tubulin, dystonin and Debrin-1 proteins were expressed at stage 6 (S6) of human endocrine differentiation (SC-endocrine) (Figure 5G and S5G). Together, these data uncover the molecular programs underlying cytoskeletal rearrangements in endocrine precursors that correlate with Syt13 expression and possibly link to its function during endocrine cell egression.

**Figure 5.**
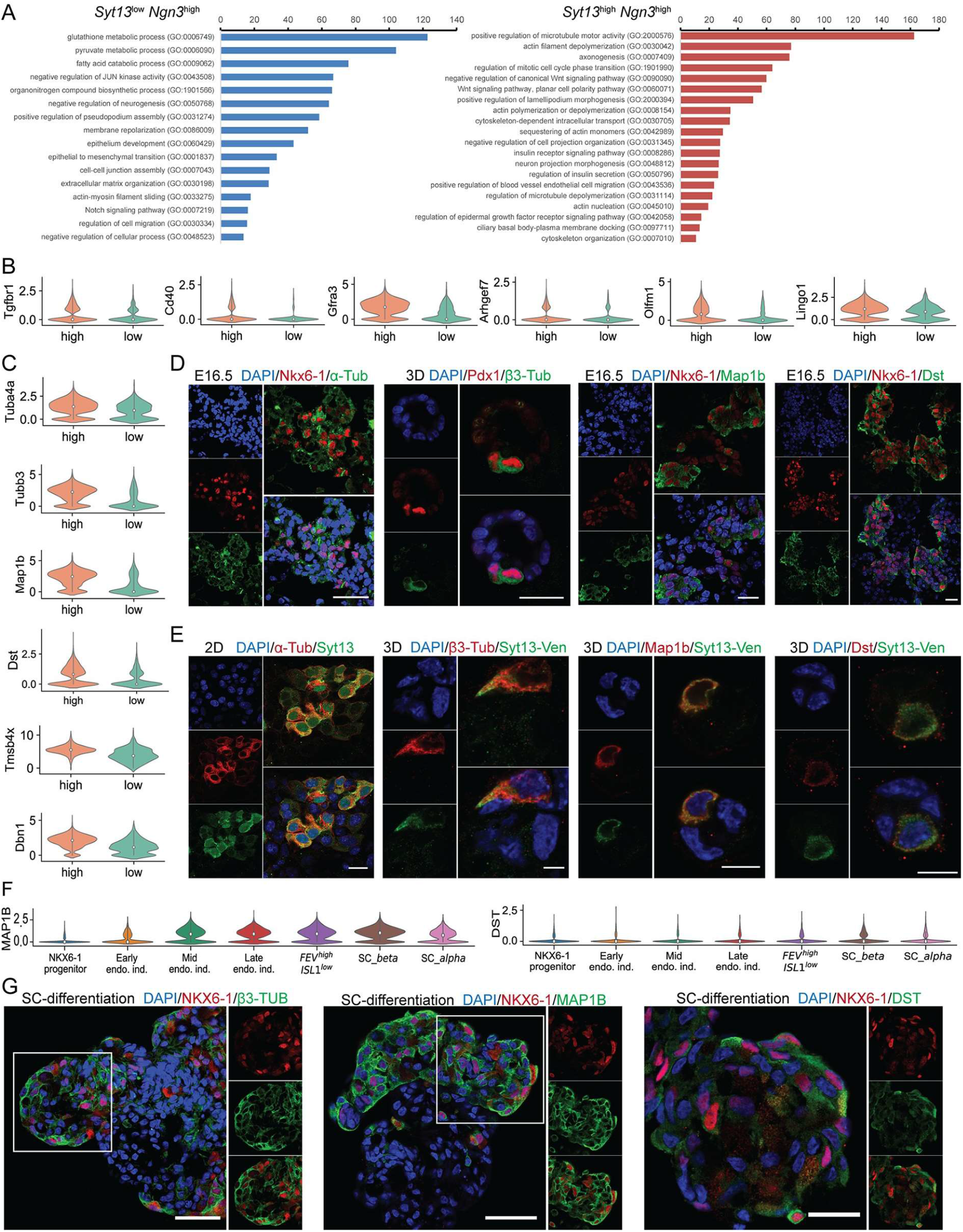
Increased Syt13 expression in endocrine precursors associates with upregulation of unique cytoskeletal components. (A) Selected terms from the pathway analysis of 500 differentially expressed genes in *Syt13*^high^ and *Syt13*^low/-^ precursors. (B) Violin plots of the expression levels of several genes involved in cell migration in *Syt13*^high^ and *Syt13*^low/-^ precursors. (C) Violin plots of the expression levels of several cytoskeletal genes in *Syt13*^high^ and *Syt13*^low/-^ precursors. (D) Immunostaining analysis indicates increased expression of α-tubulin, β3-tubulin, Map1b and Dst during endocrinogenesis. Scale bar 40 µm (α-tubulin); 20 µm (β3-tubulin, Map1b, Dst). (E) IF analysis shows co-expression of Syt13 with α-tubulin, β3-tubulin, Map1b and Dst in endocrine lineage. Scale bar 20 µm (α-tubulin); 10 µm (β3-tubulin, Map1b, Dst). (F) Violin plots of the expression levels of MAP1B and DST during human endocrinogenesis *in vitro*. (G) Immunostaining of sections derived from *in vitro* differentiated human endocrine aggregates. Scale bar 50 µm (β3-tubulin, MAP1B); 20 µm (DST). See also Figure S5.

### Syt13 interacts with the acetylated-tubulin, Map1b and dystonin cytoskeletal proteins

To explore which of the identified molecular components in the endocrine precursor and lineage cells are interlinked with Syt13 function, we performed two complementary interactome analyses. Mass spectrometry identified 79 and 69 proteins as potential Syt13 interaction partners using affinity purification and BioID proximity labeling, respectively (Figure 6A, B and Table S3). Pathway analysis revealed terms associated with cytoskeleton, vesicle trafficking, protein internalization and degradation, cell adhesion and movement (Figure S6A and S6B). Importantly, we identified α-tubulin (Tuba1a) in the direct interactome and dystonin (Dst) in both interactome lists. We confirmed the interaction of Syt13 with α-tubulin that was not hampered by deletion of Syt13 C2A or C2B domains (Figure 6C and S6C, D). α-tubulin is the major form of tubulins that can be acetylated (Ac-tub) to produce stable long-lived MTs. IF analysis disclosed the increased Ac-tub content concomitant with increased Syt13 protein levels during endocrinogenesis (Figure 6D and S6E). Moreover, the transcripts of enzymes involved in α-tubulin acetylation (*Atat1*) and deacetylation (*Hdac6* and *Sirt2*) were increased during endocrinogenesis (Figure 6E). Remarkably, these genes were not differentially expressed in *Syt13*^high^ and *Syt13*^low/-^ precursors (Figure S6F), and Ac-tub was still detected in Syt13 KO cells (Figure S6G). Yet, Syt13 was accumulated in the intracellular area enriched for Ac-tub and directly associated with this tubulin in epithelial cell lines (Figure 6F-H and S6H).

**Figure 6.**
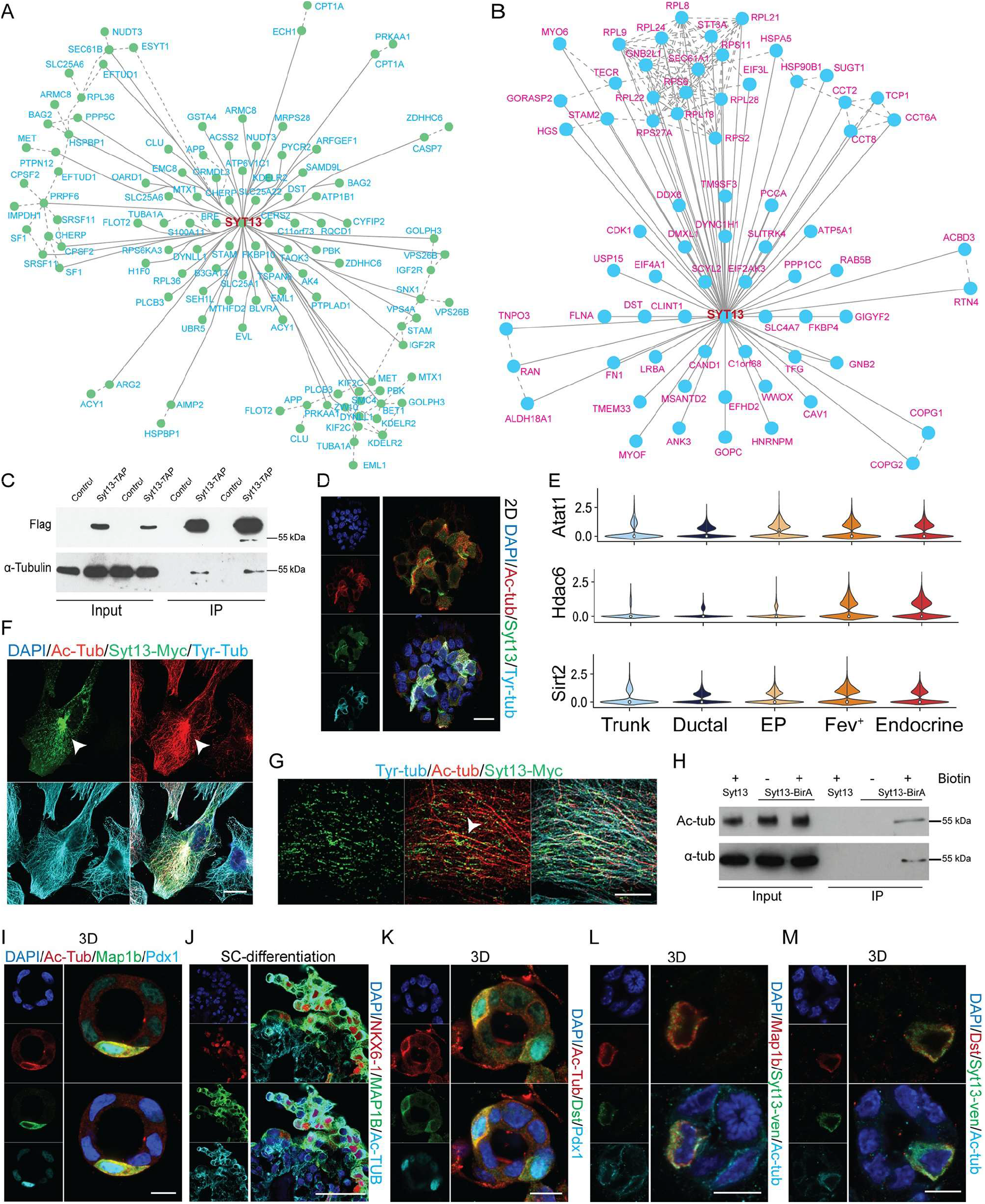
Syt13 interacts with the Ac-tub/Map1b/dystonin protein complex. (A) Direct Syt13 interacting partner proteins in MDCK cells. (B) Syt13 interacting partner proteins identified by BioID proximity labeling in MDCK cells. (C) Western blot showing Syt13 interaction with α-tubulin in MDCK cells. (D) IF analysis showing the co-expression of Syt13 and Ac-tub in endocrine cells. (E) Violin plots of the expression of enzymes involved in α-tubulin acetylation and deacetylation during endocrinogenesis. (F) Syt13 and Ac-tub accumulate at similar intracellular domains (arrowhead) in pancreatic ductal adenocarcinoma cell line (PDAC). (G) High magnification of confocal imaging of Syt13 colocalization (arrowhead) with Ac-tub in PDAC cells. (H) Western blot analysis shows close proximity of Syt13 and Ac-tub in MDCK cells. (I, J) Immunostaining analyses show the co-expression of MAP1B and Ac-TUB in mouse and human endocrine cells. (K) IF indicates co-expression of Ac-tub and dystonin in mouse endocrine cells. (L, M) Similar cellular localization of Map1b and dystonin with Syt13 protein in endocrine cells. Scale bar 10 µm (G, I, K, L, M); 20 µm (D, F); 50 µm (J). See also Figure S6 and S7.

The interconnection between Ac-tub, Map1b and dystonin and the direct interaction between Map1b and dystonin have been previously shown ^25–27^. We further found co-expression of Ac-tub with both Map1b and dystonin in mouse and human endocrine cells (Figure 6I-K and S7A). Furthermore, we revealed that both Map1b and dystonin localized at similar cellular domains to Syt13 in endocrine cells (Figure 6L and 6M). Yet, no apparent alterations in the expression levels nor the localization of both proteins was observed in Syt13 KO cells (Figure S7B and S7C). Overall, these data indicate specific upregulation of Ac-tub/Map1b/dystonin protein complex during endocrinogenesis and that Syt13 interacts with this protein complex for its trafficking and localization.

### Syt13 regulates vesicle endocytosis and remodels basal lamina

Syt13 is a leading-edge protein (Figure 4) and its KO reduces endocrine cell egression (Figure 1). Moreover, Syt13 interacts with a unique cytoskeletal protein complex (Figure 6). Since Syt13 is also a member of a vesicle trafficking protein family, these findings suggest that Syt13 might be involved in leading-edge turnover of egressing endocrine cells. In support of this idea, we identified several Syt13-interacting proteins involved in endocytosis (Caveolin-1 (Cav-1), Hgs (Hrs), Stam2, Scyl2 and Snx1) and membrane and endosomal trafficking (Ank3, Dync1h1, Dynll1, Myo6, Rab5, Rack1, Tuba1a, Vps26b and Dst), (Figure 6A, B and Table S3). Among these, we confirmed the interaction of Syt13 with Cav-1, HGS and Rab5 (Figure 7A). Furthermore, Syt13 was colocalized with Cav-1 at the PM of MDCK cells, while a fraction of this protein was localized to the intracellular vesicles (Figure 7B). Finally, we confirmed the interaction of Syt13 with Rab7 and Rab11 in IP and its colocalization with Lamp1 and Rab7 in IF, revealing the engagement of Syt13 in protein endocytosis, trafficking, degradation and recycling (Figure 7A, C and S7D).

**Figure 7.**
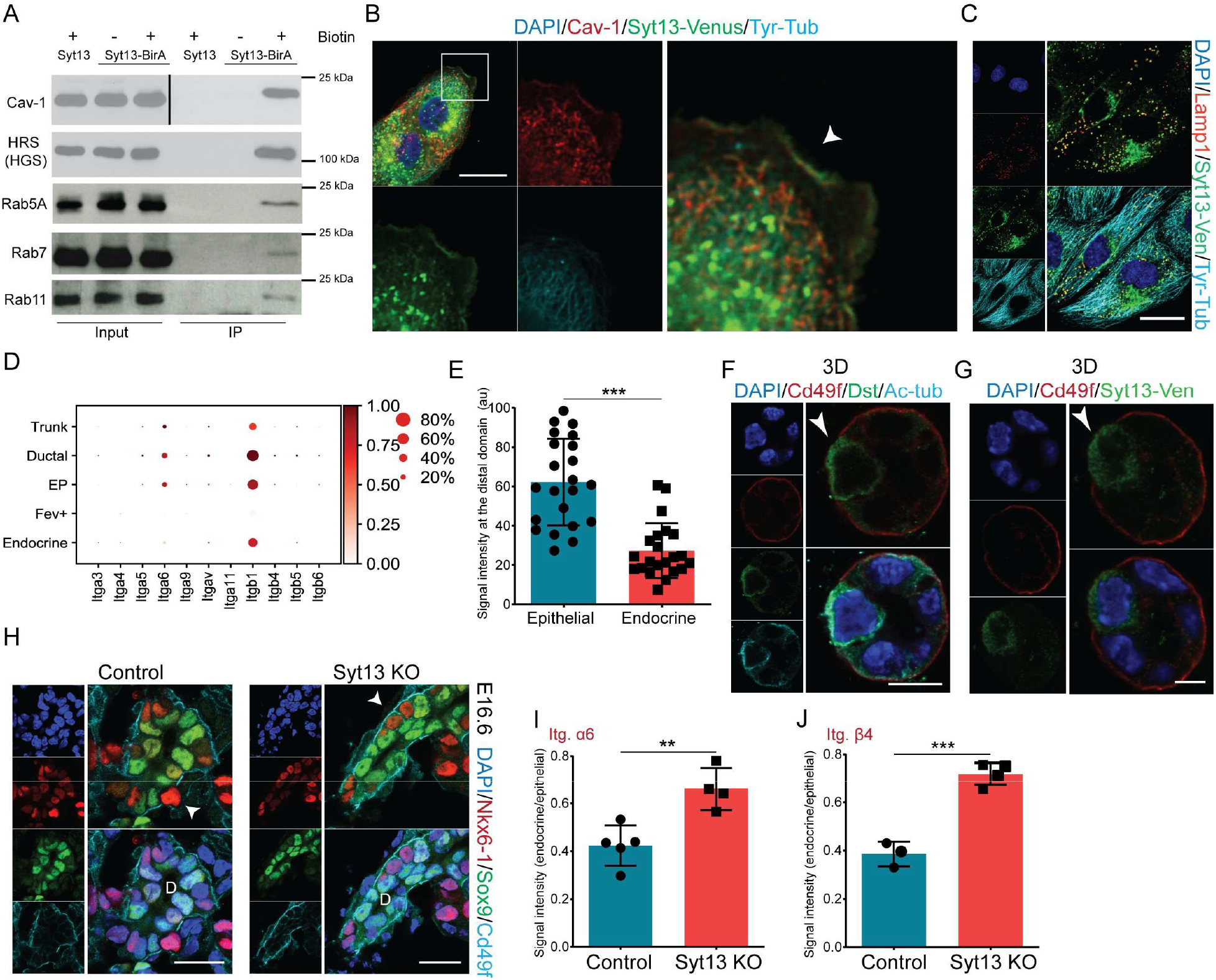
Syt13 regulates endocytosis and its knockout reduces α6β4 elimination at the endocrine leading-edge. (A) Western blot analysis indicates the close proximity of Syt13 with proteins involved in endocytosis and vesicle trafficking in MDCK cells. (B) Syt13 colocalization with Cav-1 in MDCK cells. Arrowhead indicates colocalization at the PM. (C) Colocalization of Syt13 and Lamp1 in MDCK cells. (D) Dot plot shows the expression levels of different integrin subunits during mouse endocrinogenesis. (E) Quantification of α6 integrin levels at the basal domain of embryonic epithelial and endocrine cells in 3D culture. (F, G) Reduction of α6 integrin levels in the front domain of endocrine cells (arrowhead) coincides with the expression of dystonin and Syt13 at this domain. (H) IHC analysis shows retention of α6 integrin at the basal domain (arrowheads) of Syt13 KO endocrine cells. (I, J) Quantification of α6 and β4 integrins at the basal domain of endocrine cells compared to the nearby epithelial cells. Scale bar 5 µm (G); 10 µm (F); 20 µm (B, C, H). (**P < 0.01; ***P < 0.001; t-test). Data are represented as mean ± SD. See also Figure S7.

The endocytosis function of Syt13 and its localization at the leading-edge (basal domain of precursors before or upon cell repolarization) of egressing endocrine cells, suggested its possible function in removal or recycling of the integrin-based cell-matrix adhesion complexes. To find out which integrin subunits are expressed during endocrinogenesis, we analyzed the scRNA-seq data indicating *α3, α5, α6, α9, αV, β1, β4, β5* and *β6* with different expression levels in epithelial cells (Figure 7D). Most of these integrin subunits were downregulated at EP and *Fev*^+^ states (Figure 7D), suggesting the necessity of their remodeling during endocrine cell egression. Among these, an increased expression of α6 integrin (CD49f) in acinar cells compared to lower levels in ductal and endocrine lineages has been previously reported ^28^. Having this confirmed, we further detected a reduction in the levels of CD49f in endocrine clusters compared to the ductal epithelial cells (Figure S7E). Notably, the levels of CD49f at the basal domain of the endocrine cells residing within the epithelium were reduced compared to the nearby epithelial cells (Figure 7E), indicating that the onset of α6 integrin reduction proceeds the egression process. This reduction coincided with the expression of dystonin, Syt13 and Map1b at the front domain of endocrine cells (Figure 7F, G and S7F), suggesting the possible involvement of these proteins. In support of this, we found increased levels of α6 integrin at the front domain of Syt13 KO endocrine cells compared to the control cells (Figure 7H and 7I). Similar increased levels were also observed for β4 integrin subunit (Figure 7J). These data demonstrate a function of Syt13 in the elimination of integrin-based adhesion structures at the leading-edge of egressing endocrine cells.

## Discussion

Here we present the detailed cellular and molecular underpinnings of endocrine cell egression. Setting our findings in the context of previous studies, we propose a model, in which endocrine cell egression and migration occur at EP and Fev^+^ cell states. The onset of Ngn3 expression in EPs triggers a partial EMT program ^8,15^ and results in the expression of a set of unique cytoskeletal and trafficking components, such as the polarity regulator and morphogenetic driver Syt13. Beginning at this stage, endocrine lineages condense their apical PM ^6^ marked by Syt13 protein, gradually remodel adherens junctions and reduces the apical-basal polarity components ^29^. The relocalization of Syt13 from the apical side to the leading-edge switches the apical-basal polarity to a front-rear axis essential for EPs to leave the ductal epithelium. This process is likely mediated by intracellular vesicle trafficking and requires a stable MT network consisting of proteins, such as acetylated and β3-tubulin. At the rear part of the egressing cells, the apical domain continuously narrows, which results in a formation of a tether structure connecting the cells with the epithelial plane ^7^. At the front domain, Syt13 remodels the basal lamina and modulates cell-matrix adhesion. Syt13 in concert with membrane protrusion proteins possibly generate an active and dynamic leading-edge that after abscission of the rear tether structure ^7^ enforce the detachment of the egressing cells from the epithelium.

The identity of the cellular processes, which coordinate endocrine cell egression is still a matter of debate. We found a previously unknown cell repolarization process during endocrine cell egression. Prior studies have shown reduction of the apical domain that ultimately leads to loss of apical-basal polarity in endocrine cells ^6,29^. Here, we provide evidence that after loss of apical-basal polarity, egressing and migrating endocrine cells acquire a front-rear polarity. Importantly, we found Syt13 as the first protein marking this repolarization event. The direct interaction of Syt13 with phosphoinositide phospholipids indicates the involvement of these lipids in apical-basal to front-rear repolarization. Therefore, it is likely that Syt13 is recruited to the apical or leading-edge domain through its direct interactions with PIP2 and PIP3, respectively, as it has been shown for Syt1 in neurons ^30^. Cell repolarization and the expression of actin-based membrane protrusion components indicate the existence of an active leading-edge, driving required forces for endocrine egression and directed cell migration toward the proto-islets. Additionally, we found that the ECM components are not degraded but likely remodeled during endocrine cell egression. Therefore, the newly differentiated cells that leave the epithelium join the nearby clusters with whom they share the basement membrane resulting in a local islet formation as described before ^9^.

A set of unique cytoskeleton-related genes were differentially expressed between *Syt13*^high^ and *Syt13*^low/-^ precursors including several actin-remodeling proteins. Among these were actin depolymerizing and sequestering genes such as *Tmsb4x, Scin, Cfl1* and *Mical2*, and several genes involved in actin capping, bundling and elongation such as *Dbn1, Vasp, Marcks* and *Vil1*. Along this line, the remodeling of actin cytoskeleton has been shown to regulate endocrine differentiation ^31,32^. Our analysis uncovered the molecular factors that mediate actin network rearrangements for endocrine egression. Further, we identified the increased expression of a set of particular tubulins and MT-associated proteins in *Syt13*^high^ precursors including Ac-tub, Map1b and dystonin. Strikingly, Syt13 directly interacted with Ac-tub, suggesting the dependency of Syt13 to these stable MTs for its intracellular trafficking and leading-edge localization. Moreover, the association of Syt13 with Map1b and dystonin indicates the requirement of a unique cytoskeletal railroad for Syt13-mediated vesicle trafficking. This notion together with the restricted expression of Syt13 in certain cell types, suggests that Syt13 mediates internalization and trafficking of a subset of specific membrane proteins.

Deletion of Syt13 impaired endocrine cell egression and islet morphogenesis. Furthermore, lack of this protein resulted in a shift in β- to α-cell fate. It is possible that the absence of Syt13 impacts endocrine cell egression, which consequently influences their fate decision. Along this line, lack of p120ctn in EPs has resulted in faster egression that subsequently increases their differentiation into α-cells ^33^. Contrary to this, our findings show that the delay in cell egression is linked with increased EP specification towards α-cells. Therefore, activation of a β-cell program might require a precise timing of EP occupancy within the epithelium and that faster or slower egression process induce α-cell fate. Alternatively, lack of Syt13 might directly affect β-cell differentiation. This notion is supported by the correlation between increased Syt13 expression levels in endocrine precursors and *Fev*^+^ cells with β-cell programs. As Syt13 is involved in endocytosis, it may trigger β-cell programs through controlling the involved signaling pathways and TF networks by modulating the surface receptors. Furthermore, *Syt13*^high^ precursors expressed higher levels of F-actin depolymerizing and sequestering genes. Two previous studies have shown the positive impact of F-actin network reduction on β-cell differentiation ^31,32^. Thus, it will be interesting to explore in the future whether Syt13 is directly involved in modulating actin cytoskeleton for β-cell fate decision.

We uncovered an unexpected role for the atypical Syt13 protein as endocrine cell polarity regulator and morphogenetic driver of progenitor egression. Different to the typical Syt members such as Syt1 and Syt2, Syt13 was dominantly localized to the PM of egressing endocrine cells and regulated endocytosis. A similar PM localization pattern and endocytosis function for the Syt3 in neurons have been shown ^34,35^, suggesting that Syt13 function resembles Syt3 function. Yet, the postnatal lethality of Syt13 KO mice indicates no redundant function for this protein by other Syt family members. Furthermore, together with Syt1, Syt13 deletion results in the earliest lethal phenotype among all Syt members, highlighting the unique and critical role of this protein for organ formation and function. In support of this, Syt13 is expressed in the brain ^36^, and its associated cytoskeletal proteins are also expressed in neuronal cells such as sensory neurons ^37^. In addition, a protective function of Syt13 in motor neurons of patients with neurological disorders has been shown ^38^. Thus, our findings will help to dissect the molecular action of Syt13 during neurogenesis and neurological disorders. Remarkably, SYT13 is also upregulated in several cancer cell types and its inhibition reduces cancer cell metastasis ^39–41^. These studies support a polarity and morphogenetic function of Syt13 and highlight the importance of our findings in the context of organ formation but also cancer cell dissemination.

## Methods

### Mouse lines

Mouse keeping was done at the central facilities at HMGU in accordance with the German animal welfare legislation and acknowledged guidelines of the Society of Laboratory Animals (GV-SOLAS) and of the Federation of Laboratory Animal Science Associations (FELASA). Post-mortem examination of organs was not subjected to regulatory authorization. Syt13 gene-trapped *(Syt13*^*GT/GT*^) (EUCOMM) embryonic stem cells were aggregated with CD1 morula to generate chimeric mice. Gene trap mice were bred on a mixed background. For generation of Syt13 full KO mice, first gene trap Syt13 mice were crossed with Flpe mice to obtain floxed mice *(Syt13*^*F/F*^). Then, Syt13 floxed mice were crossed with *Rosa26*^*Cre/+*^ to delete the critical exon 2 and generate the flox deleted (FD) *(Syt13*^*FD/FD*^) mice. Heterozygous *(Syt13*^*+/FD*^) intercross mice were used to obtain full knockout (KO) (Syt13 KO) embryos, which were genotyped by PCR analysis (Table S1). To generate Syt13 tissue-specific conditional knockout mice, *Syt13*^*F/F*^ mice were crossed with constitutive *Tg (Neurog3-cre)C1Able/J (Ngn3*^*+/Cre*^) ^42^ and *Ins1*^*Cre*^ *(Ins1*^*tm1(cre)Thor*^*) (Ins1*^*+/Cre*^) mice ^43^. Syt13-Venus fusion mouse line, in which endogenous Syt13 is fused with the fluorescent protein Venus has been recently generated by us and will be described elsewhere.

### Cloning, cell culture and transfection

Cloning was performed using standard protocols. Different Syt13 constructs were generated using the pCAG mammalian expression vector (Table S1). Cell transfection was performed using lipofectamine™ 2000 transfection reagent (Gibco/Invitrogen GmbH, 11668019). For generating stable cell lines, cells were transfected with linearized constructs and were treated with 1-2 µg/ml puromycin for 5 days and the surviving mixed clones were expanded.

MDCK cells (NBL-2) (Sigma, 85011435) and pancreatic ductal adenocarcinoma (PDAC) cell lines ^44^ were cultured and maintained in a standard medium (DMEM, 10% FBS, penicillin/streptomycin). For 2D stainings, cells were trypsinized and then plated on µ-Slide 8-well chambers (Ibidi, 80826) followed by the experimental procedure. For 3D culture, µ-Slide 8-well chambers (Ibidi, 80821) were coated with 100% grocontrolh factor-reduced Matrigel (BD Biosciences) for 15 min at 37 °C. Single MDCK cells were resuspended in 2% Matrigel-containing medium. MDCK cysts were fixed and stained after 3-4 days.

Primary pancreatic epithelial cyst culture was performed as described previously ^29^. Briefly, E14.5 mouse embryonic pancreata were dissected and kept in 0.25% trypsin-EDTA for 15-30 min on ice and then 5 min at 37 °C. After replacing the trypsin with culture medium and 30 times pipetting up and down, single-cell suspension was achieved. Single pancreatic cells were then cultured on 100% Matrigel-coated µ-Slide 8-well chambers (Ibidi, 80821) in the culture medium containing 2% Matrigel. The PECs were fixed and stained after 1-2 days.

2D pancreatic culture was established by dissecting E14.5 embryonic pancreata and incubation in 0.25% trypsin-EDTA for 15-30 min on ice followed by 5 min incubation at 37 °C. The trypsin was replaced by culture medium and partial digestion was performed by pipetting the samples up and down for 10 times. Cell clusters were cultured on ˂10% Matrigel-coated µ-Slide 8-well chambers (Ibidi, 80821) in the previously described condition medium ^29^.

Pancreatic explant culture was performed by dissecting E13.5 pancreata into 3-4 pieces and culturing them in µ-Slide 8-well chambers (Ibidi, 80821) coated with 50 µg/mL bovine fibronectin (Sigma, F1141). Samples were treated by 14 µM PIP3/protein binding inhibitor (PIT-1) (Abcam, ab120885) or 10 µM LY294002 (Promega, V1201). The medium was replaced with a fresh one every 24 h. Samples were fixed and stained after 2 days in culture.

The differentiation of human iPSCs into pancreatic endocrine cells *in vitro* was performed using previously described protocol ^45^ with slight modifications ^29^.

### Immunostaining and imaging

Dissected embryonic pancreata or explant culture samples were fixed in 4% PFA in PBS for 2 h overnight at 4 °C. The tissues were merged in 10% and 30% sucrose-PBS solutions at RT (2 h each solution) followed by 1:1 solution 30% sucrose:tissue-freezing medium (Leica 14020108926). Afterwards, they were embedded in cryoblocks using tissue-freezing medium and sections of 20 μm thickness were cut using a cryostat. Next, the samples were permeabilized (0.1% Triton, 0.1 M Glycine) for 30 min and incubated in blocking solution (10% FCS, 3% Donkey serum, 0.1% BSA and 0.1% Tween-20 in PBS) for 1 h at room temperature (RT). Then, the primary antibodies (Table S1) diluted in the blocking solution were added to the samples overnight at 4 °C. After washing with PBS they were stained with secondary antibodies (Table S1) diluted in the blocking solution for 3-5 h at RT. The samples were then incubated with 4’, 6-diamidin-2-phenylindol (DAPI), followed by washing with PBS and embedding in commercial medium (Life Tech., ProLong Gold).

2D and 3D cultures of cell lines or primary pancreatic cells were fixed in 4% PFA (12 min at 37 °C) followed by 10 min permeabilization (100 mM Glycine and 0.2% Triton X-100) at RT. After 3x washing, cells were incubated with blocking solution for 30 min at RT and then incubated with primary antibodies (Table S1) for 1-3 h at RT. After washing, cells were incubated with secondary antibodies (Table S1) for 1 h at RT and DAPI was added and samples were embedded.

All images were obtained with a Leica microscope of the type DMI 6000 using the LAS AF software. Images were analyzed using LAS AF and ImageJ software programs.

### Cell sorting and quantitative PCR (qPCR) analysis

Embryonic pancreata from NVF; Syt13 KO at E15.5 were dissected. Next, individual pancreata were kept in 0.25% Trypsin for 5 min on ice and then incubated at 37 °C for 10 min. The single-cell samples were then centrifuged at 1700 rpm for 5 min at 4 °C. 5 µl anti-mouse CD326 (EpCAM) PE (eBioscience, 12-5791-81) and rat IgG2a K isotype control (eBioscience, 12-4321-42) were used for 1×10^2^ cells in 100 µl total volume. After staining for 15 min at 4 °C, the cells were washed with PBS and stained with DAPI for 5 min to detect dead cells. The samples were then washed and resuspended in the FACS buffer (PBS, 1% BSA, 0.5 mM EDTA) and loaded for FACS sorting. The gating strategy was as follows: main population> single cells > living cells (DAPI negative)> EpCam^+^ (PE^+^) and Ngn3^+^ (FITC^+^) cells. The cells were collected in Qiazol (Qiagen, 79306). After RNA isolation, a MessageAmp™ II aRNA Amplification Kit (Thermo Fisher Scientific) was utilized to maximize the yield per sample. The procedure for the amplification was undertaken as stated in the kit’s protocol. Briefly, as first step a cDNA amplification synthesis by reverse transcription was prepared, followed by cDNA purification. Next, an *in vitro* transcription reaction was prepared to generate multiple copies of amplified RNA (aRNA) from the double-stranded cDNA templates providing aRNA. Finally, the aRNA was purified and quantified. qPCR analysis was done using TaqMan™ probes (Life Technologies; Table S1) and 25 ng cDNA per reaction. Each reaction consisted of 4.5 µL cDNA in nuclease-free water, 5 µL TaqMan™ Advanced master mix (Life Technologies) and 0.5 µL TaqMan probe™. The qPCR was performed using Viia7 (Thermo Fisher Scientific). Ct-values were normalized among samples, transformed to linear expression values, normalized on reference genes and on control samples.

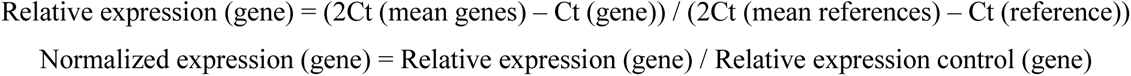

### Western blotting

Protein samples were loaded on SDS-PAGE gel and then were transferred to Nitrocellulose membrane (Bio-Rad, Cat. Nr. 1620112). After blocking for 1 h at RT, membranes were incubated with the primary antibodies at 4 °C overnight (Table S1) followed by 3x washing and then incubating with the secondary antibodies (Table S1) for 1 h at RT. Next, membranes were incubated with Pierce ECL Western Blotting Substrate (Thermo Fisher Scientific) or SuperSignal West Pico PLUS Chemilumineszenz-Substrat (Thermo Fisher Scientific) and the signals were detected by enhanced chemiluminescence.

### Immunoprecipitation using strep-tag (affinity purification)

1 × 10^2^ MDCK cells stably expressing recombinant Syt13 protein variants fused with TAP-tag were cultured in 10 cm^2^ dishes in control medium for 3 days at 37 °C. Cells were washed with 10 ml ice-cold PBS and were lysed with 700 µl lysis buffer consisting of 1x TBS (Tris-buffered saline), 0.5% (v/v) Igepal CA 630, protease and phosphatase inhibitors. Samples were scraped and kept on ice for 15 min followed by centrifugation for 10 minutes at 21470 × g at 4 °C. The supernatant was collected and a fraction of it was stored at -20 °C as Input. The remaining supernatant was added to 50 µl G-Protein Sepharose beads (Protein G-Sepharose 4 Fast Flow, GE Healthcare) (pre-washed with the lysis buffer) and were kept rotating at 4 °C for 1 h to capture the proteins that nonspecifically interact with the Sepharose beads. After centrifugation the samples were incubated with 50 µl Strep-Tactin Sepharose beads (Superflow 50% suspension, IBA) (pre-washed 3x with the lysis buffer) for 1 h at 4 °C while rotating. After centrifugation the supernatants were discarded and the beads were washed 3x with the washing buffer (1 × TBS, 0.1% (v/v) Igepal CA 630 and phosphatase inhibitor). The samples were then processed for western blotting or mass spectrometry as follows.

For western blotting, the beads were resuspended in the sample buffer, heated for 5 min at 95 °C. After cooling down on ice the samples were centrifuged for 10 min at 21470 × g and the supernatants were collected as the IP sample, which together with the input samples were applied to western blotting.

For mass spectrometry the beads were incubated with 150 µL the elution buffer (GE Healthcare, 2-1042-025) for 10 min at 4 °C while rotating. Samples were centrifuged for 30 second at 1000 × g at 4 °C. The supernatant was collected and stored at -80 for further processing.

### Immunoprecipitation using BioID approach

1 × 10^2^ MDCK cells stably expressing Syt13 or Syt13-BirA recombinant proteins were cultured in 14 cm^2^ dishes in grocontrolh medium for 3 days at 37 °C. The medium was replaced with a fresh grocontrolh medium containing 50 µM biotin and after 16 h each dish was lysed with 1 ml lysis buffer (50 mM Tris buffer pH 7.5, 0.5% Sodiumdesoxycolate, 150 mM NaCl, 1% NP40, 0.1% SDS) and kept on ice. In parallel, 240 µL Pierce™ High Capacity NeutrAvidin™ Agarose beads (ThermoFisher Scientific, Cat: 29202) were washed 3x with the lysis buffer and resuspended in 1 mL lysis buffer. Next, the cell lysates were centrifuged for 30 min at 14000 rpm at 4 °C and 3 mL supernatant for each condition was collected. 90 µL of the samples were stored as the input samples and the rest were mixed with 300 µL beads-lysis buffer in the 15 mL falcon tubes rotating for 3 h at RT. Next, samples were centrifuged for 5 min at 500 rpm at 4 °C. Supernatants were discarded and the beads were used for western blotting or mass spectrometry as follows. For western blotting, the beads were washed 3x with the washing buffer (50 mM Tris buffer pH 7.5, 0.5% Sodiumdesoxycolate, 150 mM NaCl, 1% NP40, 0.5% SDS) and 2x with 1x TBS. The beads were then resuspended in 90 µL sample buffer, heated for 5 min at 95 °C. After cooling down on ice the samples were centrifuged and the supernatants were collected as the IP sample, which together with the input samples were applied to western blotting.

For mass spectrometry, 500 µl TBS was added to the beads and mixed rigorously. After centrifugation at 7000 × g for 30 second supernatants were removed and the washing was repeated two more times. 60 µl elution buffer (2 M urea, 50 mM Tris-HCl pH 7.5 and 5 μg/mL Trypsin (SIGMA; T6567-5×20UG)) (twice as the bead volume) was added and samples were incubated for 1 h at 27 °C under constant agitation (800 rpm in a thermo mixer) followed by centrifugation. Supernatants were collected in a fresh Eppendorf tube. The beads were then resuspended with 60 µl of the elution buffer (containing 2 M urea, 50 mM Tris-HCl pH 7.5 and 1 mM DTT), centrifuged and the supernatant was collected and pooled with the previous one. This last step was repeated and 180 µl total volume per each reaction was obtained. Samples were left at RT to continue to digest overnight and were stored at -80 °C for further mass spectrometry procedure.

### Mass spectrometry

Affinity purified eluates were precipitated with chloroform and methanol followed by trypsin digestion as described before ^46^. LC-MS/MS analysis was performed on Ultimate3000 nanoRSLC systems (Thermo Scientific) coupled to an Orbitrap Fusion Tribrid mass spectrometer (Thermo Scientific) by a nanospray ion source. Tryptic peptide mixtures were injected automatically and loaded at a flow rate of 30 μl/min in 0.1% trifluoroacetic acid in HPLC-grade water onto a nano trap column (300 μm i.d. × 5 mm Pre column, packed with Acclaim PepMap100 C18, 5 μm, 100 Å; Thermo Scientific). After 3 minutes, peptides were eluted and separated on the analytical column (75 μm i.d. × 25 cm, Acclaim PepMap RSLC C18, 2 μm, 100 Å; Thermo Scientific) by a linear gradient from 2% to 30% of buffer B (80% acetonitrile and 0.08% formic acid in HPLC-grade water) in buffer A (2% acetonitrile and 0.1% formic acid in HPLC-grade water) at a flow rate of 300 nl/min over 117 minutes. Remaining peptides were eluted by a short gradient from 30% to 95% buffer B in 5 minutes. Analysis of the eluted peptides was done on an LTQ Fusion mass spectrometer. From the high-resolution MS pre-scan with a mass range of 335 to 1500, the most intense peptide ions were selected for fragment analysis in the orbitrap depending by using a high speed method if they were at least doubly charged. The normalized collision energy for HCD was set to a value of 27 and the resulting fragments were detected with a resolution of 120,000. The lock mass option was activated; the background signal with a mass of 445.12003 was used as lock mass ^47^. Every ion selected for fragmentation was excluded for 20 seconds by dynamic exclusion. MS/MS data were analyzed using the MaxQuant software (version 1.6.1.0) ^48,49^. As a digesting enzyme, Trypsin/P was selected with maximal 2 missed cleavages. Cysteine carbamidomethylation was set for fixed modifications, and oxidation of methionine and N-terminal acetylation were specified as variable modifications. The data were analyzed by label-free quantification with the minimum ratio count of 3. The first search peptide tolerance was set to 20, the main search peptide tolerance to 4.5 ppm and the re-quantify option was selected. For peptide and protein identification the human subset of the SwissProt database (release 2014_04) was used and contaminants were detected using the MaxQuant contaminant search. A minimum peptide number of 2 and a minimum length of 7 amino acids was tolerated. Unique and razor peptides were used for quantification. The match between run option was enabled with a match time window of 0.7 min and an alignment time window of 20 min. The statistical analysis including ratio, t-test and significance A calculation was done using the Perseus software (version 1.6.2.3) ^50^.

Interaction candidates from mass spectrometry were used to gain insights into the biological processes involved with Syt13. We used Metascape ^51^ to perform the pathway enrichment analysis.

### Surface biotinylation

1 × 10^2^ MDCK cells stably expressing Syt13-TAP recombinant proteins were cultured in 6-well plates in grocontrolh medium for 3 days at 37 °C. The cells were washed 3x with ice-cold PBS pH 8.0. Cells were then incubated with 500 µL of 2 mM EZ-link Sulfo-NHS-Biotin (Thermo Scientific, 21425) in PBS for 15 min on ice. Samples were then washe 3x with PBS containing 100 mM glycine and were lysed with lysis buffer (1x TBS, 0.5% NP40, protease inhibitor) for 15 min on ice. After centrifugation, 30 µL of supernatants were stored as input and the rest were incubated with the lysis buffer-prewashed G-sepharose beads (50 µL of 50% slurry per each reaction; Protein G-Sepharose 4 Fast Flow, GE Healthcare) for 3 h at 4 °C while rotating. After centrifugation, the supernatants were incubated with lysis-buffer prewashed high capacity NeutrAvidin agarose resin beads (50 µL of 50% slurry per each reaction) (Thermo Scientific, 29202) overnight at 4 °C rotating. The beads were then washed 3x with the washing buffer (1x TBS, 0.1% NP40) and 40 µL sample buffer were added. Together with the input fractions, the samples were applied to western blotting.

### Purification of recombinant domains of Syt13 protein variants

Three different constructs expressing recombinant Syt13 protein variants fused with Myc-tag were generated using the pGEX-6P-1 vector. These included Syt13-C2AB (aa 39 to 426 of mouse Syt13), Syt13-C2A (aa 39 to 282 of mouse Syt13), Syt13-C2B (aa 289 to 426 of mouse Syt13). Recombinant protein production was carried out in Escherichia coli BL21 (DE3). Bacterial cells were grown in a total of 700 ml LB medium supplemented with ampicillin (100 µg/ml) to reach an OD A600 of 0.7 at 37 °C, induced with 0.5 mM isopropyl β-D-1-thiogalactopyranoside, and incubated for 3-6 h at 25 °C. Bacterial lysate was prepared by sonication in lysis buffer (50 mM Tris pH 8.0, 300 mM NaCl, 4 mM DTT, 2 mM EDTA), followed by three passes through a Emulsiflex C3 homogenizer (Avestin Inc., Ottawa, Canada). After removal of cell debris by centrifugation for 15 min at 20 000 × g at 4 °C, the supernatant was incubated with 5 ml Glutathione Sepharose 4B (GE Healthcare) for 1 h at 4 °C in batch mode under gentle rotation. Beads were packed into the column, washed with 20 column volumes (CV) wash buffer 1 (50 mM Tris pH 8.0, 750 mM NaCl, 4 mM DTT, 2 mM EDTA, 50 mM KCL, 10 mM MgCl2, 1 mM ATP) and subsequently with 20 CV wash buffer 2 (50 mM Tris pH 8.0, 750 mM NaCl, 2 mM DTT). Beads were then incubated with recombinant GST-tagged human rhino-virus 3C protease (50 µg per CV) for 16-18 h at 4 °C and the protein was eluted with 6 CV elution buffer (50 mM Tris pH 8.0, 150 mM NaCl, 1 mM DTT). The eluted protein was concentrated using ultrafiltration units with 10 kDa regenerated cellulose membrane (Amicon Ultra, Merck Millipore) and subsequently loaded on an equilibrated (50 mM Tris pH 8.0, 150 mM NaCl, 1 mM DTT, 5% glycerol) Superdex 75 10/300 GL column (GE Healthcare) for size exclusion chromatography. Protein purity was assessed by SDS-PAGE (NuPAGE gels; 12% bis-tris gels, and MOPS buffer both Invitrogen). Dynamic Light Scattering (Zetasizer nano ZS, Malvern Instruments, Worcestershire, UK) was used to confirm monomeric state. For protein quantification Bradford assay1 or BCA protein assay kit (Pierce, Thermo Fisher Scientific) was used.

### Preparation of large unilamellar vesicles (LUVs)

*1*-*palmitoyl*-*2*-*oleoyl phosphatidylcholine* (POPC) and phosphoinositides were purchased from Avanti Polar Lipids, Alabaster; cholesterol was purchased from Sigma-Aldrich, Germany. POPC and chloroform stocks were dissolved in chloroform, phosphoinositides in chloroform:methanol:water 1:2:0.8. The fatty acid distribution consists of C18:1, mimicking unsaturated species found within organisms. Lipid concentration and stability of the stocks was validated by a total phosphorus assay ^52^ and thin layer chromatography (TLC) at regular basis. For vesicle preparation, lipids were mixed in desired mol % ratio (POPC/cholesterol/phosphoinositide 65/30/5 mol %, POPC/cholesterol 70/30 mol %) and dried under nitrogen gas stream, followed by incubation under vacuum for 2-16 h to remove organic solvents. The dried lipid film was re-hydrated in liposome buffer (10 mM Hepes, 150 mM NaCl, pH 7.4) to a final concentration of 1 mg/ml, for 15 min at 300 rpm. Multilamellar vesicles were then subjected to 10 cycles of freezing in liquid nitrogen and subsequent thawing in a heating block at 18 °C. The vesicle solution was extruded 21 times through a 100 nm diameter polycarbonate membrane (Whatman® Nuclepore, Fisher Scientific) using an extrusion kit (Avanti Polar Lipids, Alabaster). For quality control, vesicles were subjected to dynamic light scattering (DLS) and zeta potential measurements (see following section). Lipid composition and stability of vesicles was analyzed by TLC.

### Size and zeta potential measurements

Size and zeta potential of Large Unilamellar Vesicles (LUV), and protein size were determined using a Zetasizer Nano ZS (Malvern Instruments, UK). For size measurements LUVs were diluted to a final concentration of lipids ranging from 0.08 to 0.4 mg/ml. Size of proteins was measured at a concentration above 0.5 mg/ml in buffer (50 mM Tris, 150 mM NaCl, 1 mM DTT, pH 8.0). Importantly, all buffers were filtered (PVDF, 0.45 µm, Millipore) prior to use. Zeta-potential of LUVs was measured at a total lipid concentration of 0.08 mg/ml in filtered (PVDF, 0.45 µm, Millipore) water.

### Thin layer chromatography

TLC was used to validate lipid composition of lipid stocks and LUVs. LUVs were extracted with choloroform:methanol:aceton:1N HCl (2:1:0,5:0,1 v/v), where the organic phase was dried under nitrogen stream and dissolved in 20 μl of extraction solution. Silica-coated TLC plates (HPTLC Silica Gel 60, 10 × 10 cm, Merck Darmstadt) were pre-treated with 1% potassium oxalate in methanol:water (2:3 v/v) for 12-16 h and dried under vacuum at RT. The extracts were spotted on TLC plates and placed in a closed glass chamber with chloroform:acetone:methanol:acetic acid:water (46:17:15:14:8 v/v). Subsequently, the TLC plate was air-dried and lipids were visualized by spraying the plate with primuline (Sigma-Aldrich) solution (5% primuline in acetone:water 8:2 v/v), which was then scanned on a Typhoon 9410 imager using 457 nm laser (GE Healthcare).

### Electrochemiluminescence-based immunoassay

The binding assay was performed using Meso Scale Discovery 384 well high bind plates as described previously ^53^. In brief, all binding experiments were carried out at 22 °C. Liposomes (2 μl) were passively adsorbed on the electrode surface for 1 h, and residual sites on the surface were blocked for 1 h with 0.25% porcine gelatin (Sigma-Aldrich) in TRIS buffer (50 mM Tris, 150 mM NaCl, pH 8.0). After three washing steps with TRIS buffer, serial dilutions of recombinant protein in blocking buffer was added to the respective wells and incubated for 2 h. Unbound protein was removed, and anti-c-myc antibody (clone 9E10, Santa Cruz Biotechnology) at 1.25 µg/ml concentration in 0.25% porcine gelatin was applied for 1 h followed by three subsequent wash steps with TRIS buffer. For detection, a secondary anti-mouse antibodies labeled with Sulfo-TAG (Meso Scale Discovery) was used at 1.25 μg/ml in blocking buffer for 1 h in the dark. Free secondary antibody was washed off, and reading buffer (surfactant-free reading buffer from Meso Scale Discovery) was added. The readout was performed on a Meso Scale Discovery SECTOR Imager 6000 chemiluminescence reader. Data were analyzed with GraphPad Prism 6.07. First, signal from PC/cholesterol (70/30 mol %) vesicles was subtracted as a background. Next, a non-linear curve fitting was applied, and the binding kinetics were calculated using one site – specific binding fitting.

### ScRNA-seq data sources and analysis

For all scRNA-seq data analyses python 3.7.6 and the Scanpy package (v1.4.4) (https://github.com/theislab/scanpy) were used ^54^. Processed and normalized scRNA-seq data and cell annotations of mouse embryonic pancreatic cells ^21^ were downloaded from GEO (accession number: GSE132188). The mouse data include cells from four embryonic stages (E12.5-E15.5) from the pancreatic epithelium and was enriched for endocrine cells by using an Ngn3-reporter. To classify cells as *Syt13*^high^ and *Syt13*^low^ we applied a threshold of 1 to *Syt13* expression values (normalized counts). To infer lineage relationships between *Syt13*^high^ and *Syt13*^low^ precursors and *Fev*^+^ and hormone^+^ clusters partition-based graph abstraction (PAGA) ^55^ was performed as implemented in Scanpy *tl*.*paga* with a threshold of 0.05 for cluster connectivity. As input to PAGA, we recomputed the single-cell nearest neighbor-graph (kNN) on the 50 first principal components computed on top highly variable genes (*pp*.*highly_variable* with default parameters) and a local neighborhood size of 15. For differential expression testing a t-test as implemented in tl.rank_genes_groups was used. Genes expressed in <20 cells of the subset of cells used for testing were excluded. For GO term enrichment the top ranked genes sorted based on the t-test statistic were used (top 500 for EP clusters, top 300 for FEV+ clusters). GO term enrichment was performed with the gseapy (v0.10.2) implementation of EnrichR ^56^.

Processed and normalized scRNA-seq data and cell annotations of human *in vitro* stem cell differentiation ^22^ were downloaded from GEO (accession number GSE114412). The human data contains cells sampled from stage 5 of the differentiation protocol.

Raw scRNA-seq data from human fetal pancreas were downloaded from the data visualization center descartes (https://descartes.brotmanbaty.org/bbi/human-gene-expression-during-development/) ^23^, and loaded into R to convert the rds-file to an AnnData object for downstream analysis with the rpy2 (v3.3.5, https://github.com/rpy2/rpy2) and anndata2ri (v1.0.4, https://github.com/theislab/anndata2ri) python packages. Raw counts were normalized using total count normalization and log-transformed (log(count+1)). For the per cell normalization factor highly expressed genes in a cell were not considered (i.d. the parameter *exclude_highly_expressed* was set to *True* in *pp*.*normalize_total*). To identify endocrine lineage populations we iteratively clustered and annotated cells with the louvain clustering method as implemented in the louvain-igraph package (v0.7.0, https://github.com/vtraag/louvain-igraph) ^57^ and adopted by Scanpy in *tl*.*louvain* (for details see available analysis notebook). For each round of clustering a single-cell kNN using 15 neighbors was computed on the 50 first principal components of the expression matrix of the 4000 top highly variable genes. Genes expressed in <10 cells were excluded before normalization and clustering. Clusters were annotated based on marker genes and merged if expressing the same markers. Mesenchymal clusters were distinguished from epithelial cells (endocrine lineage) based on the expression of the markers *EPCAM* and *VIM*, and excluded. *FEV*+ endocrine precursors were annotated using *FEV* as a marker. *NEUROG3* was not detected. Endocrine lineage precursors were annotated based on co-expression of *FEV* and the transcription factors *ARX* for α-cell precursors and *PAX4* for β-cell precursors. Hormone+ clusters were annotated with the hormones *GCG* for α-cells, *INS* for the β-cells, *SST* for δ-cells, and *GHRL* for Ɛ-cells.

### Code availability

Custom notebooks for all analyses of scRNA-seq data will be made available in a github repository upon publication.

### Statistical analysis

All statistical analysis was performed on GraphPad prism 9 and presented in figure legends.

### Image quantification

#### Endocrine egression in vivo

Quantification was performed using confocal images from randomly selected pancreas areas. The quantification was done by counting the ratio (as percentage) of Nkx6-1^high^/Sox9^-^ cells that were residing within or in the direct contact with the epithelium to the total Sox9^+^ cells. For each condition, more than 11 confocal images from randomly selected pancreas areas from 4 animals were used. The analysis was conducted using LAS-AF software.

#### Endocrine cluster direct attachment to the epithelium

Quantification was performed using confocal images from randomly selected pancreas areas. We divided the area of epithelium-proto islets direct attachment to the total area of proto-islet periphery Areas were detected via “wand tracing tool”, supervised by adjusting the tolerance level based on the quality of the staining. The surface of contact was manually drawn. The analysis was conducted using Fiji ImageJ.

#### Ngn3 quantification

Quantification of the endocrine progenitor (Ngn3^+^) cells was performed on confocal images from randomly selected pancreas areas by using automatic nuclei counts with IMARIS software (Bitplane). Different developmental stages (E13.5-E15.5) were analyzed from 3 controls and 3 Syt13 KO mice for each stage. Each animal value was generated by the average of n ≥ 4 pancreatic sections with a minimum number of 5000 DAPI cells per mice.

#### α-/β-cell ratio quantification

For the quantification of α- to β-cell ratio, we used randomly selected pancreatic sections where hormone-expressing cells were manually counted using the Leica LAS-AF software. Full Syt13 KO and 2 different CKOs (*Ngn3*^*+/Cre*^*;Syt13*^*F/FD*^ and *Ins1*^*+/Cre*^; *Syt13*^*F/FD*^) were analyzed with n≥ 3 mice for each condition. Each mice value consists in an average of minimum 4 pancreatic sections.

#### Syt13 apical localization

Signal intensity of the recombinant Syt13 protein variants at the apical domain were divided to the total cytosolic signal. 32 total epithelial cysts from each condition were quantified from 4 independent experiments. The analysis was conducted using LAS-AF software.

#### Endocrine delamination in the explant culture

Quantification was performed using confocal images from randomly selected pancreas areas. The quantification was done by counting the percentage of GFP^+^ cells that were residing within the epithelium of the total epithelial cells. For each condition, more than 65 epithelial domains from 3 independent experiments were quantified. The analysis was conducted using LAS-AF software.

#### α6 and β4 integrin subunit

Quantification was performed using confocal images from randomly selected pancreas areas. The signal intensity of the respective proteins at the basal domain of endocrine cells was divided to the signal at the basal domain of epithelial cells. For each protein, n ≥ 20 images from n ≥ 3 independent experiment were quantified. The analysis was conducted using LAS-AF software.

## Acknowledgements

We thank Anika Boettcher, Ralph Boettcher and Mara Catani for the helpful comments and discussion. We thank Michael Sterr, Bianca Vogel, Anne Savoca, Maximilian Schuster, Mahfuza Akter, Johanna Tüshaus, Kerstin Diemer, Robert Fimmen and Ines Kunze for technical support. The work was supported by the Helmholtz Society, Helmholtz Portfolio Theme ‘Metabolic Dysfunction and Common Disease, German Research Foundation, and German Center for Diabetes Research (DZD e.V.) for financial support. This project was supported by DZD, DZD-Next funding (M.B., Ü.C. and H.L.) and the Tistou & Charlotte Kerstan Stiftung (M.U. and K.B.).

## Author contribution

Conceptualization, M.B. and H.L.; Methodology, M.B., A.B.P., E.N. and K.B.; Formal Analysis, S.T.; Investigation, M.B., A.B.P., M.T.M., E.N., K.S., J.J., P.C., C.S., K.B., S.J.W., N.H. and I.B.; Writing – Original Draft, M.B.; Writing – Review & Editing, A.B.P. and H.L.; Supervision, M.B., F.J.T., Ü.C. and H.L.; Funding Acquisition, M.B., K.B., M.U., Ü.C. and H.L. Further information and requests for resources and reagents should be directed to and will be fulfilled by the lead contact Heiko Lickert.

The authors declare no competing interests.

---

**Supplementary Figure 1.**
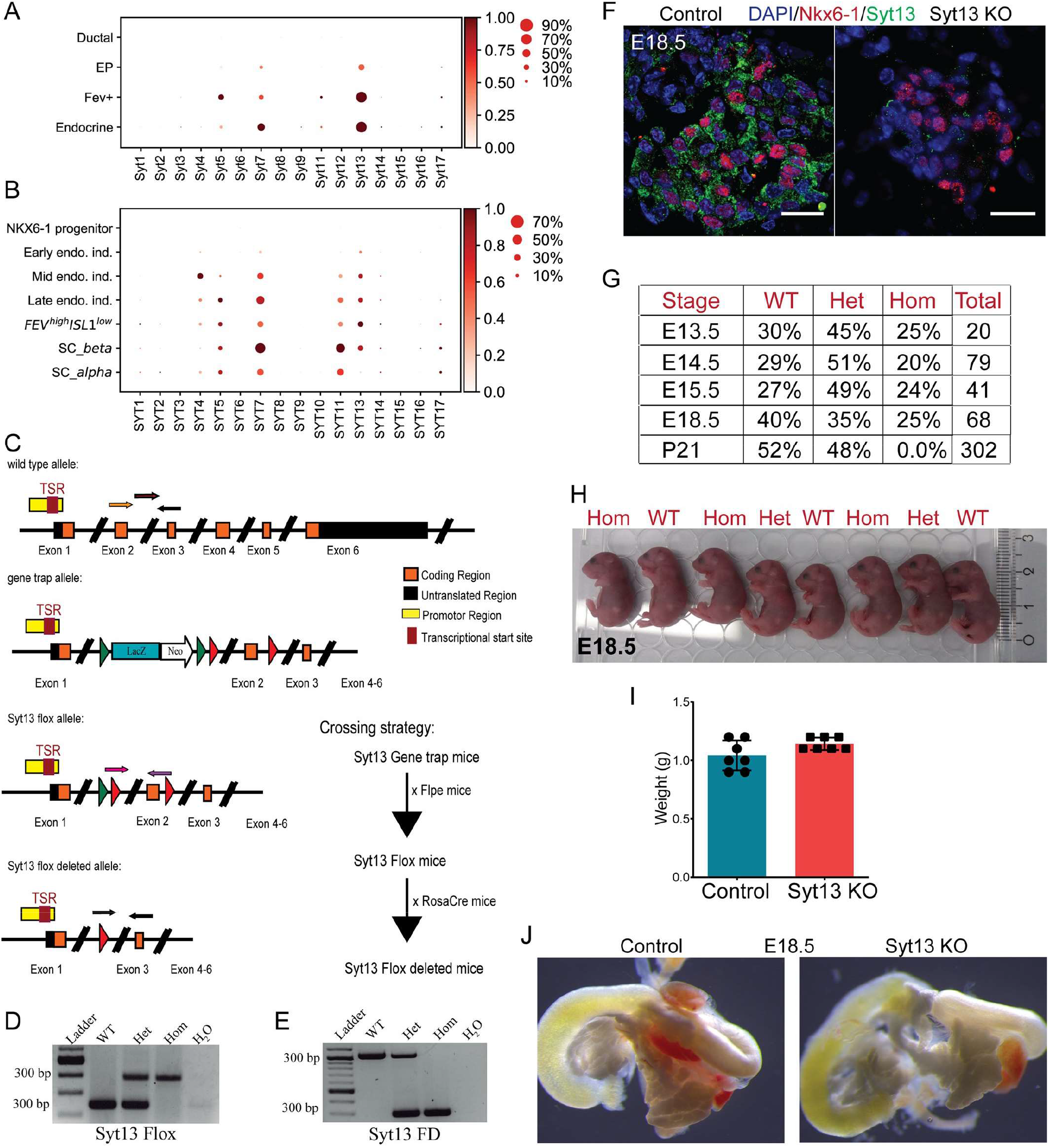
Generation and characterization of Syt13 KO mice. (A, B) Dot plots showing the expression of Syt family members during mouse (A) and human in vitro (B) endocrinogenesis in scRNA-seq data. (C) Schematic representation of Syt13 allele and the strategy for generation Syt13 Flox-deleted (FD) allele to generate *Syt13*^*FD/FD*^ (Syt13 knockout (Syt13 KO)) mice. (D, E) PCR analyses for genotyping *Syt13*^*Flox*^ and full-body *Syt13*^*FD/FD*^ mice. (F) IHC of pancreatic sections confirms deletion of Syt13 protein in Syt13 KO mice. Scale bar 20 µm. (G) Mendelian ratio of offspring resulting from the cross between heterozygous *(Syt13*^*+/FD*^) mice shows no embryonic but postnatal lethality. (H) Gross morphological analysis and (I) the weight of Syt13 KO and control embryos at E18.5. (J) Gross morphological analysis of Syt13 KO and control pancreata at E18.5.

**Supplementary Figure 2.**
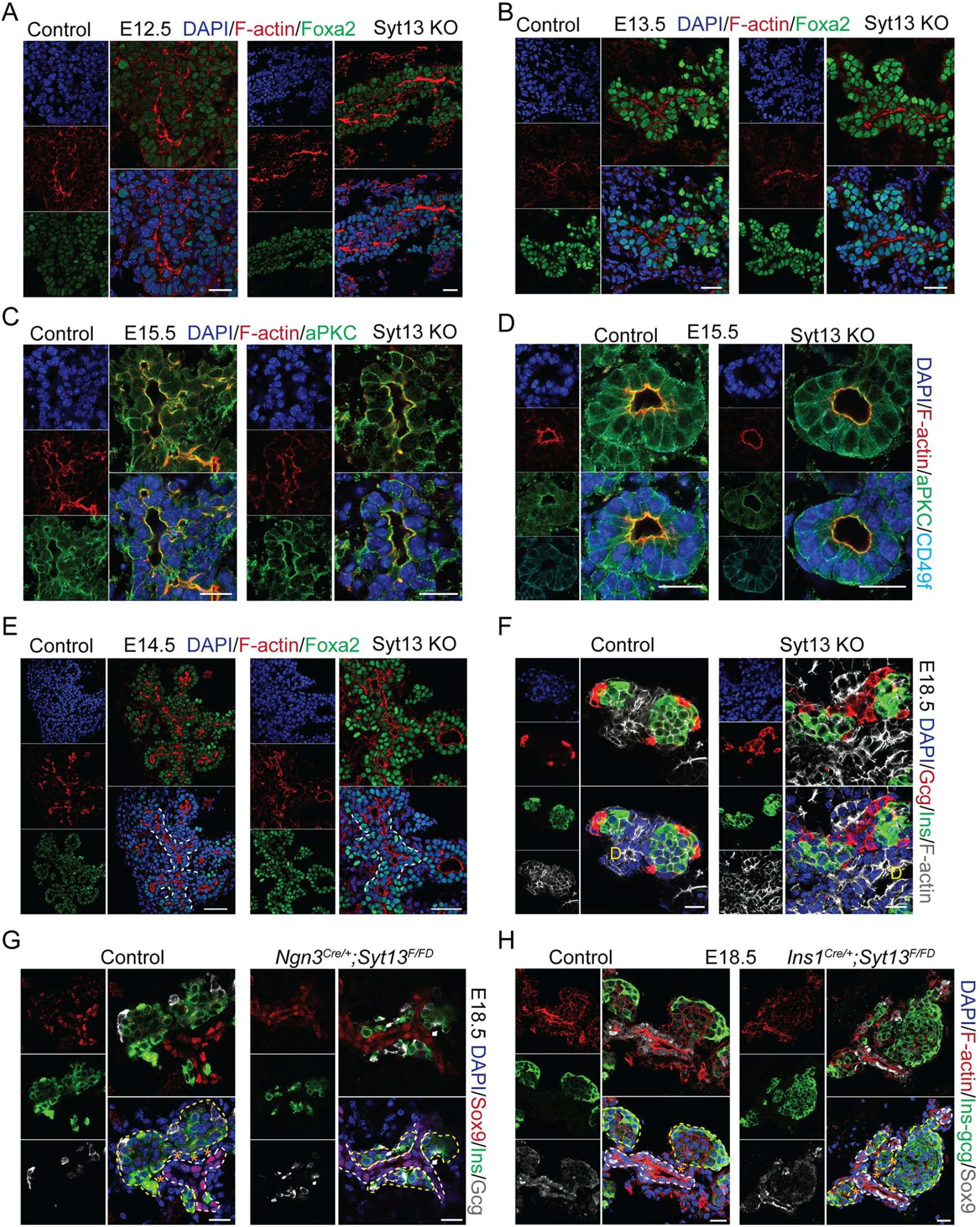
Epithelium architecture and polarity in Syt13 KO mice. (A, B) Comparable epithelial organization between control and Syt13 KO pancreata at E12.5 and E13.5. (C) Normal apical-basal polarity of ductal epithelium in control and Syt13 KO pancreata. (D) Normal apical-basal polarity of acinar cells in Syt13 KO pancreata. (E) A multi-layer epithelium appears in Syt13 KO pancreata. The extra layer is composed of Foxa2^high^ cells. (F) Different α- and β-cells arrangement and positioning within the proto-islets in Syt13 KO compared to control. (G) Endocrine contact are with nearby epithelium in *Ngn3*^*Cre/+*^*;Syt13*^*F/FD*^ compared to control pancreatic sections. (H) Endocrine contact are with nearby epithelium in *Ins1*^*Cre/+*^*;Syt13*^*F/FD*^ and control pancreatic sections. D, duct. Yellow dashed lines indicate proto-islet clusters and white dashed lines indicate epithelium. Stars indicate space between endocrine and epithelial cells. Scale bar 20 µm (A-D, F-H); 40 µm (E).

**Supplementary Figure 3.**
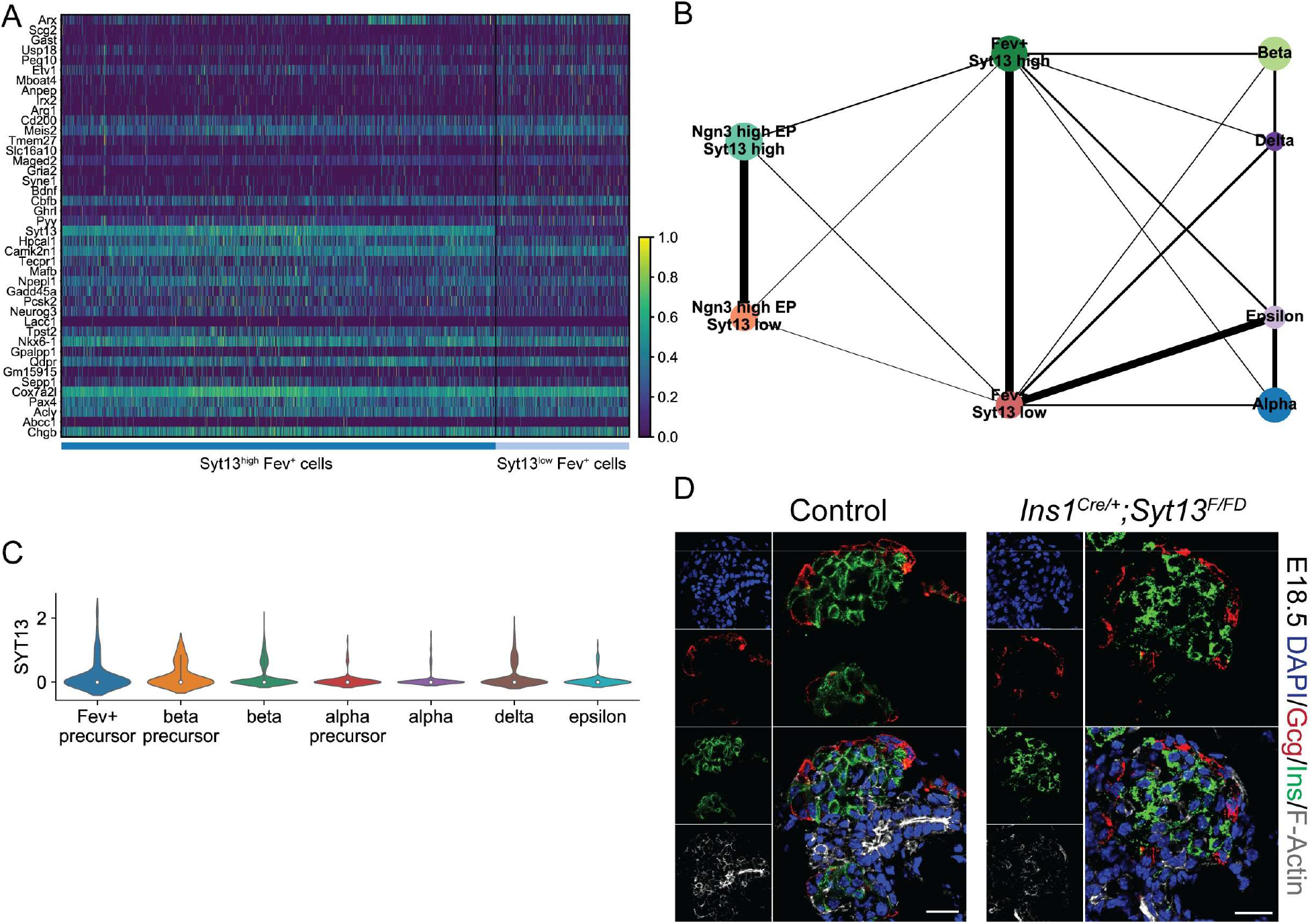
Increased levels of Syt13 in endocrine lineage link with β-cell fate. (A) Headmap of top 20 differentially expressed genes in *Syt13*^high^ and *Syt13*^low/-^ *Fev*^+^ cells. (B) PAGA analysis corroborates the lineage relationship between *Syt13*^high^ and *Syt13*^low/-^ precursors and *Fev*^+^ cells with different types of hormone^+^ endocrine cells. (C) Violin plots showing *Syt13* expression in different endocrine lineages from human fetal pancreas. (D) IHC analysis of pancreatic sections from *Ins1*^*Cre/+*^*;Syt13*^*F/FD*^ mice. Scale bar 20 µm.

**Supplementary Figure 4.**
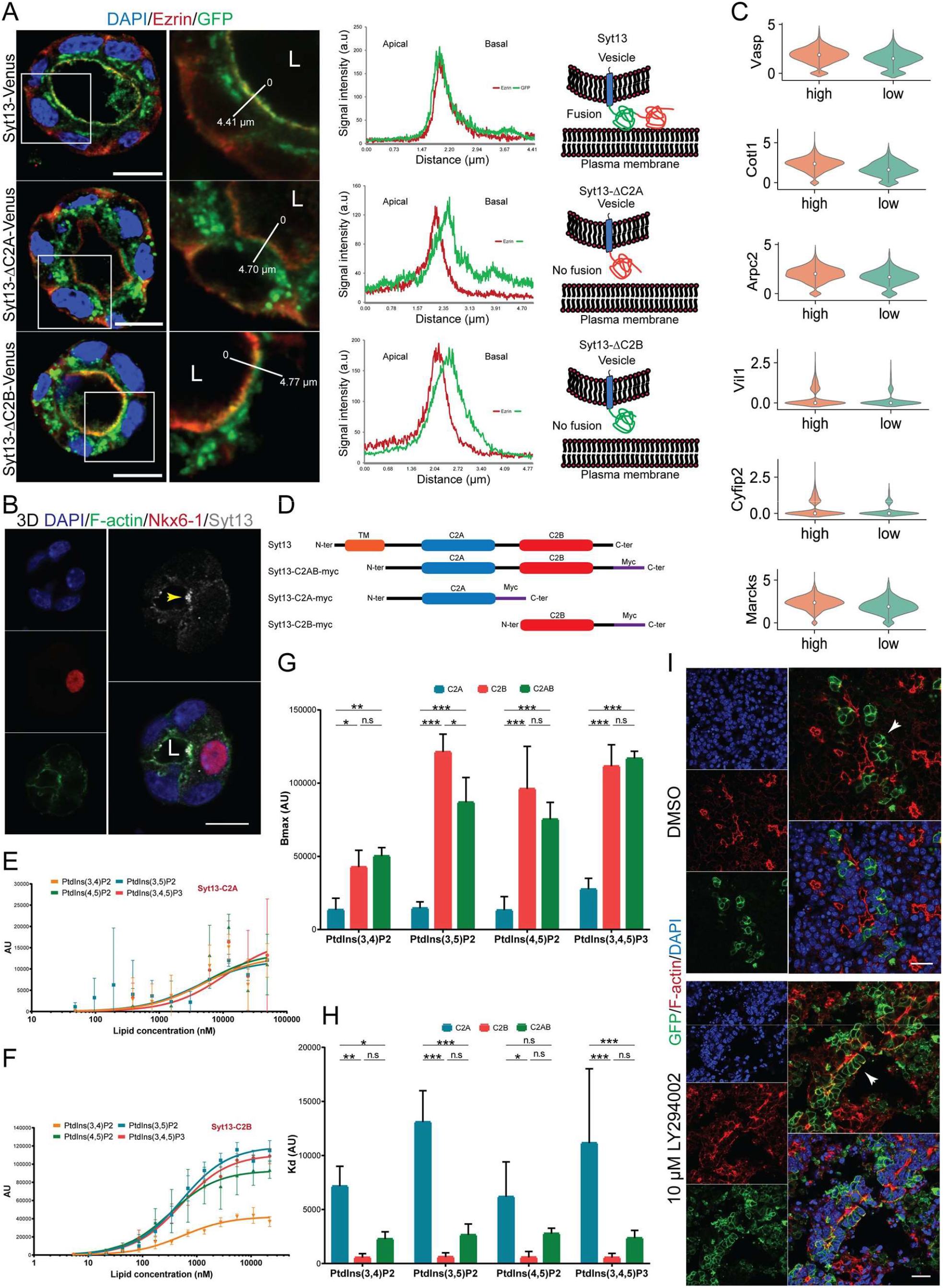
Syt13 polarized localization in epithelial cells and its lipid-binding property. (A) IF and plot profiles showing the accumulation of truncated Syt13 proteins under the apical domain in 3D MDCK epithelial cysts, demonstrating that PM docking and fusion depends on the C2A and C2B domains. (B) Syt13 protein and F-actin colocalize (arrowhead) at the apical domain of differentiated endocrine cells residing within the pancreatic epithelium. (C) Violin plots showing expression levels of several lamellipodia-related genes in *Syt13*^high^ and *Syt13*^low/-^ precursors. (D) Generation of different purified Syt13 protein variants for lipid-binding analysis. (E) Binding of purified Syt13 C2A domain to 100 nm LUVs containing POPC/cholesterol/phosphoinositide 65/30/5 mol % followed by electrochemiluminescence-based immunoassay. (F) Binding of purified Syt13 C2B domain to LUVs containing POPC/cholesterol/phosphoinositide 65/30/5 mol % followed by liposomal electrochemiluminescence-based immunoassay. (G, H) Bmax and Kd of different truncated Syt13 variants binding to LUVs containing POPC/cholesterol/phosphoinositide 65/30/5 mol %. (I) Treatment of explant culture of E13.5 *Ngn3*^*Cre/+*^; *ROSA26*^*mTmG/mTmG*^ pancreata with PI3K inhibitor LY294002 for 48 h, impairs endocrine egression. Arrowheads indicate endocrine cells. L, lumen. Scale bar 10 µm (A, B); 20 µm (J). (n.s, non-significant; *P < 0.05; **P < 0.01; ***P < 0.001; t-test). Data are represented as mean ± SD.

**Supplementary Figure 5.**
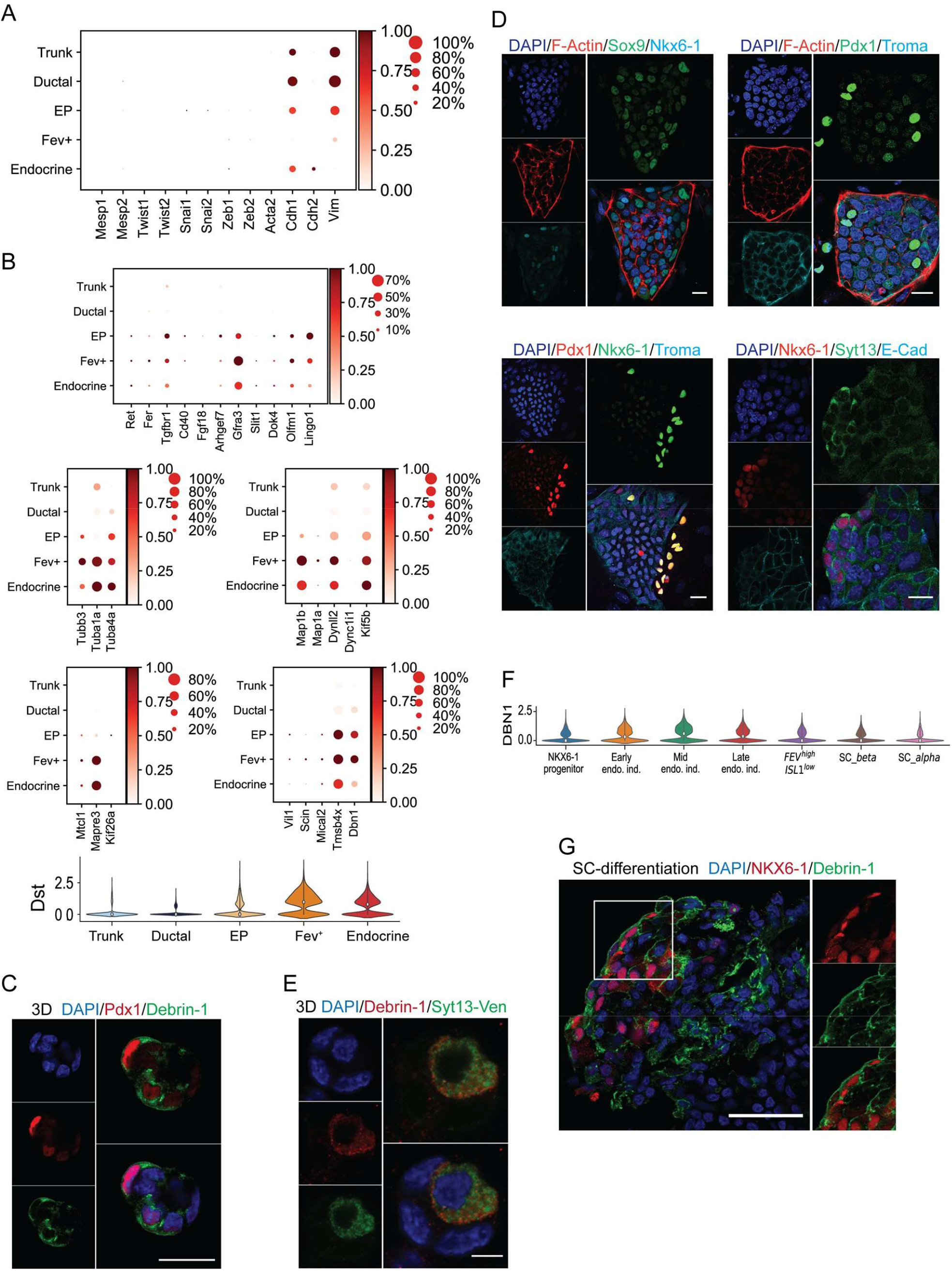
Expression dynamics of EMT and cytoskeletal related genes during endocrinogenesis. (A) Dot plot showing the expression levels of EMT-related genes during mouse endocrinogenesis. (B) Dot plots and violin plot showing the expression of several genes involved in the cell dynamics process at different stages of endocrinogenesis. (C) Characterization of the newly established 2D culture of mouse embryonic pancreatic epithelial cells. Scale bar 20 µm. (D) Increased expression of Debrin-1 during endocrinogenesis. Scale bar 20 µm. (E) IF analysis shows co-expression of Syt13 with Debrin-1 in endocrine lineage. Scale bar 5 µm. (F) Violin plots of the expression levels of *DBN1* during human endocrinogenesis *in vitro*. (G) Increased expression of Debrin-1 in human endocrine cells. Scale bar 50 µm.

**Supplementary Figure 6.**
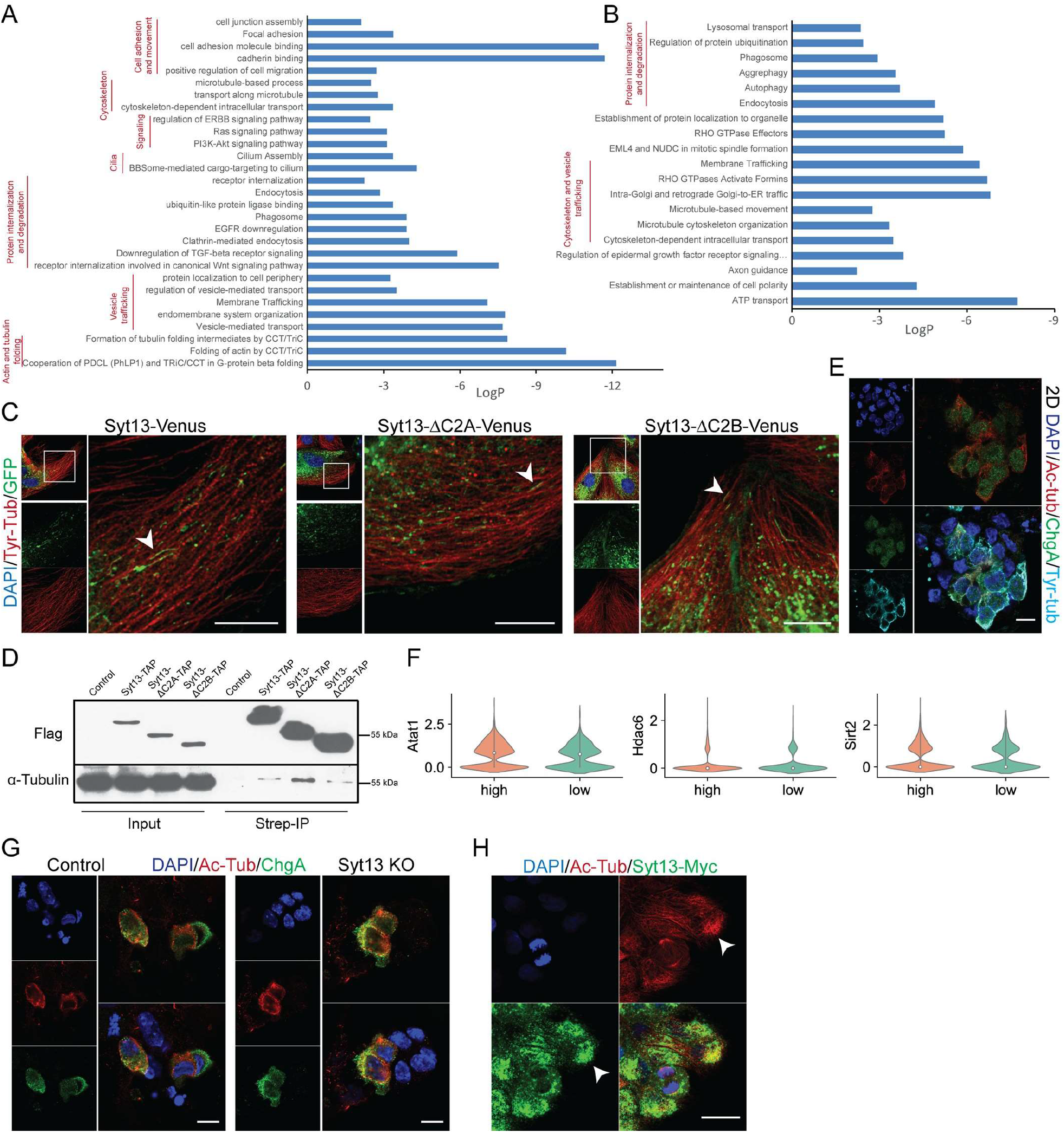
Syt13 interacts with acetylated tubulin. (A, B) Selected terms of pathway analysis of Syt13 interactome partners identified by proximity labeling approach (A) and affinity purification (B). (C) Association of Syt13 full-length protein and its truncated variants with tubulin cytoskeleton (arrowheads) in MDCK cells. (D) IP showing the interaction of Syt13 full-length protein and its truncated variants with α-tubulin in MDCK cells. (E) IF analysis discloses increased levels of acetylated-tubulin in endocrine cells in the 2D system. (F) Violin plots showing the expression levels of enzymes involved in α-tubulin acetylation and deacetylation in *Syt13*^high^ and *Syt13*^low/-^ precursors. (G) Detection of Ac-tub in endocrine cells from control and Syt13 KO endocrine cells using the 2D system. (H) IF analysis of Syt13-overexpressing MDCK cells shows enrichment of Syt13 and Ac-tub in similar intracellular domains (arrowheads). Scale bar 10 µm (C, E, G); 20 µm (H).

**Supplementary Figure 7.**
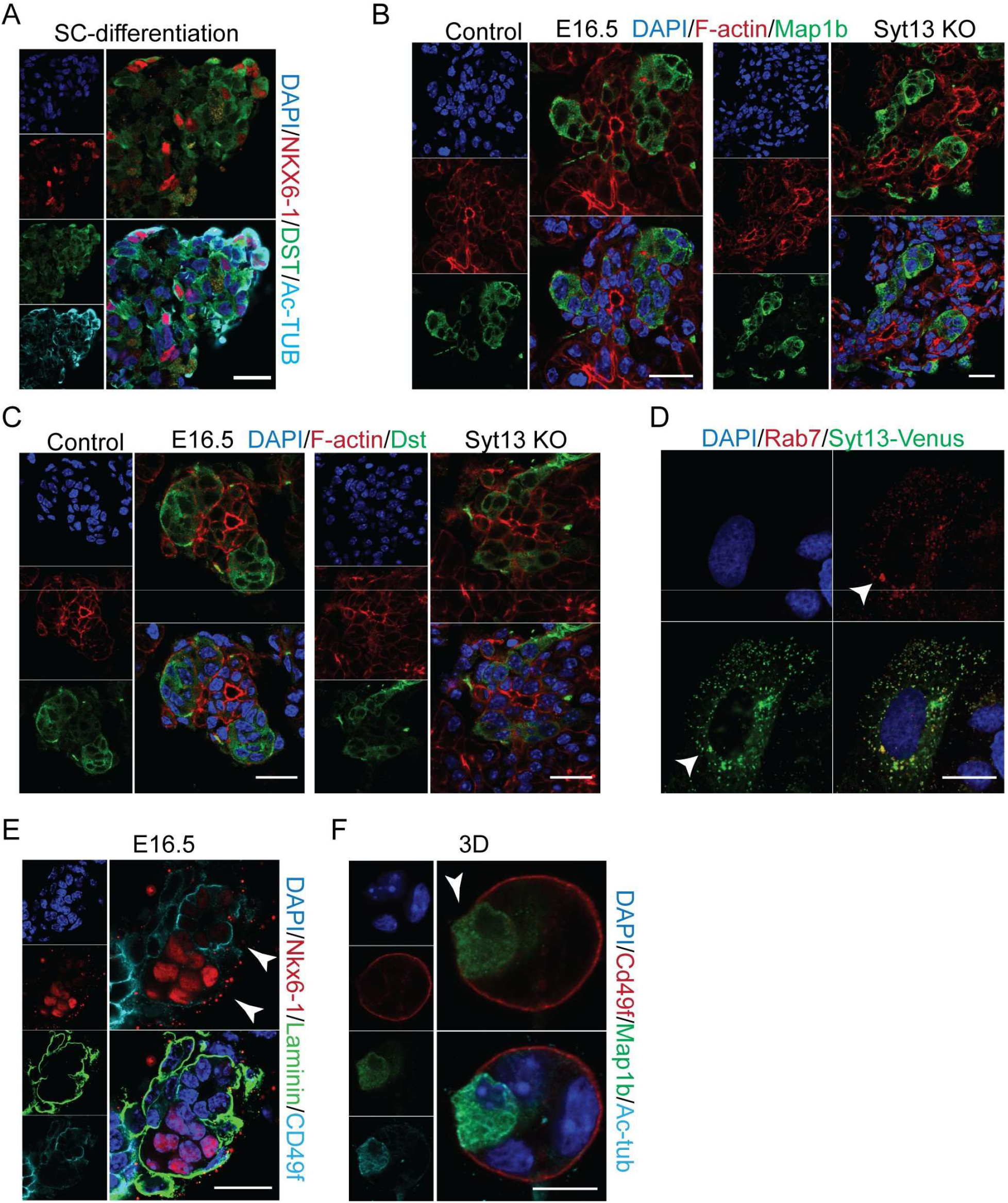
No impact of Syt13 KO on Map1b and dystonin expression. (A) Immunostaining analyses show the co-expression of dystonin and Ac-TUB in human endocrine cells. (B, C) Normal expression and localization of Map1b and dystonin in Syt13 KO and control pancreatic sections. (D) Syt13 and Rab7 colocalize (arrowhead) in MDCK cells. (E) Reduction of CD49f but not laminin in endocrine clusters compared to the ductal epithelial cells stained in pancreatic sections at E16.5. (F) Reduction of α6 integrin levels in the front domain of endocrine cells (arrowhead) coincides with the expression of Map1b at this domain. Scale bar 10 µm (F); 20 µm (A-E).

**Supplementary Table 1**. Detailed information of genotyping primers, generated plasmids, used primary and secondary antibodies and used qPCR probs.

**Supplementary Table 2**. Differential expressed genes in *Syt13*^low^ and *Syt13*^high^ endocrine precursors and *Fev*^+^ cells and the corresponding pathway analyses.

**Supplementary Table 2**. List of potential Syt13 interaction partner proteins identified by affinity purification and BioID proximity labeling and the corresponding pathway analysis.

## Notes

### Competing Interest Statement

The authors have declared no competing interest.

## References

1. Bechard, M. E. et al. Precommitment low-level neurog3 expression defines a long-lived mitotic endocrine-biased progenitor pool that drives production of endocrine-committed cells. Genes Dev. 30, 1852–1865 (2016).

2. Byrnes, L. E. et al. Lineage dynamics of murine pancreatic development at single-cell resolution. Nat. Commun. 9, 1–17 (2018).

3. Kesavan, G. et al. Cdc42-Mediated Tubulogenesis Controls Cell Specification. Cell 139, 791–801 (2009).

4. Pan, F. C. & Wright, C. Pancreas organogenesis: from bud to plexus to gland. Dev. Dyn. 240, 530–65 (2011).

5. Bastidas-Ponce, A., Scheibner, K., Lickert, H. & Bakhti, M. Cellular and molecular mechanisms coordinating pancreas development. Dev. 144, (2017).

6. Löf-Öhlin, Z. M. et al. EGFR signalling controls cellular fate and pancreatic organogenesis by regulating apicobasal polarity. Nat. Cell Biol. 19, 1313–1325 (2017).

7. Bankaitis, E. D., Bechard, M. E., Gu, G., Magnuson, M. A. & Wright, C. V. E. ROCK-nmMyoII, Notch and Neurog3 gene-dosage link epithelial morphogenesis with cell fate in the pancreatic endocrine-progenitor niche. Development 145, dev162115 (2018).

8. Gouzi, M., Kim, Y. H., Katsumoto, K., Johansson, K. & Grapin-Botton, A. Neurogenin3 initiates stepwise delamination of differentiating endocrine cells during pancreas development. Dev. Dyn. 240, 589–604 (2011).

9. Sharon, N. et al. A Peninsular Structure Coordinates Asynchronous Differentiation with Morphogenesis to Generate Pancreatic Islets. Cell 176, 790-804.e13 (2019).

10. Greiner, T. U., Kesavan, G., Ståhlberg, A. & Semb, H. Rac1 regulates pancreatic islet morphogenesis. BMC Dev. Biol. 9, 2 (2009).

11. Kesavan, G. et al. Cdc42/N-WASP signaling links actin dynamics to pancreatic β cell delamination and differentiation. Development 141, 685–96 (2014).

12. Pictet, R. & Rutter, W. J. Development of the embryonic endocrine pancreas. Handbook of Physiology 25–66 (1972).

13. Pauerstein, P. T. et al. A radial axis defined by semaphorin-to-neuropilin signaling controls pancreatic islet morphogenesis. Development 144, 3744–3754 (2017).

14. Freudenblum, J. et al. In vivo imaging of emerging endocrine cells reveals a requirement for PI3K-regulated motility in pancreatic islet morphogenesis. Development 145, dev158477 (2018).

15. Rukstalis, J. M. & Habener, J. F. Snail2, a mediator of epithelial-mesenchymal transitions, expressed in progenitor cells of the developing endocrine pancreas. Gene Expr. Patterns 7, 471–9 (2007).

16. Chapman, E. R. How does synaptotagmin trigger neurotransmitter release? Annual Review of Biochemistry vol. 77 615–41 (2008).

17. Südhof, T. C. & Rizo, J. Synaptotagmins: C2-domain proteins that regulate membrane traffic. Neuron vol. 17 379–88 (1996).

18. Fukuda, M. & Mikoshiba, K. Characterization of KIAA1427 protein as an atypical synaptotagmin (Syt XIII). Biochem. J. 354, 249–257 (2001).

19. von Poser, C. & Südhof, T. C. Synaptotagmin 13: structure and expression of a novel synaptotagmin. Eur. J. Cell Biol. 80, 41–47 (2001).

20. Willmann, S. J. et al. The global gene expression profile of the secondary transition during pancreatic development. Mech. Dev. 139, 51–64 (2016).

21. Bastidas-Ponce, A. et al. Comprehensive single cell mRNA profiling reveals a detailed roadmap for pancreatic endocrinogenesis. Development (2019) doi:10.1242/dev.173849.

22. Veres, A. et al. Charting cellular identity during human in vitro β-cell differentiation. Nature doi: 10.1038/s41586-019-1168-5. (2019) doi:10.1038/s41586-019-1168-5.

23. Cao, J. et al. A human cell atlas of fetal gene expression. Science (80-.). 370, (2020).

24. Yang, J. et al. Guidelines and definitions for research on epithelial–mesenchymal transition. Nat. Rev. Mol. Cell Biol. 21, 341–352 (2020).

25. Ryan, S. D. et al. Microtubule stability, Golgi organization, and transport flux require dystonin-a2-MAP1B interaction. J. Cell Biol. 196, 727–742 (2012).

26. Bhanot, K., Young, K. G. & Kothary, R. MAP1B and clathrin are novel interacting partners of the giant cyto-linker dystonin. J. Proteome Res. 10, 5118–5127 (2011).

27. Takemura, R. et al. Increased microtubule stability and alpha tubulin acetylation in cells transfected with microtubule-associated proteins MAP1B, MAP2 or tau. J. Cell Sci. 103, 953–964 (1992).

28. Sugiyama, T., Rodriguez, R. T., McLean, G. W. & Kim, S. K. Conserved markers of fetal pancreatic epithelium permit prospective isolation of islet progenitor cells by FACS. Proc. Natl. Acad. Sci. 104, 175–80 (2007).

29. Bakhti, M. et al. Establishment of a high-resolution 3D modeling system for studying pancreatic epithelial cell biology in vitro. Mol. Metab. 30, 16–29 (2019).

30. Martin, T. F. J. PI(4,5)P2-binding effector proteins for vesicle exocytosis. Biochim. Biophys. Acta Mol. Cell Biol. Lipids 1851, 785–793 (2015).

31. Mamidi, A. et al. Mechanosignalling via integrins directs fate decisions of pancreatic progenitors. Nature 564, 114–118 (2018).

32. Hogrebe, N. J., Augsornworawat, P., Maxwell, K. G., Velazco-Cruz, L. & Millman, J. R. Targeting the cytoskeleton to direct pancreatic differentiation of human pluripotent stem cells. Nat. Biotechnol. 38, 460–470 (2020).

33. Nyeng, P. et al. p120ctn-Mediated Organ Patterning Precedes and Determines Pancreatic Progenitor Fate. Dev. Cell 49, 31–47 (2019).

34. Butz, S., Fernandez-Chacon, R., Schmitz, F., Jahn, R. & Südhof, T. C. The subcellular localizations of atypical synaptotagmins III and VI. Synaptotagmin III is enriched in synapses and synaptic plasma membranes but not in synaptic vesicles. J. Biol. Chem. 274, 18290–18296 (1999).

35. Awasthi, A. et al. Synaptotagmin-3 drives AMPA receptor endocytosis, depression of synapse strength, and forgetting. Science (80-.). 363, (2019).

36. Han, S. et al. Altered expression of synaptotagmin 13 mRNA in adult mouse brain after contextual fear conditioning. Biochem. Biophys. Res. Commun. 425, 880–5 (2012).

37. Ferrier, A., Boyer, J. G. & Kothary, R. Cellular and Molecular Biology of Neuronal Dystonin. International Review of Cell and Molecular Biology vol. 300 (Elsevier, 2013).

38. Nizzardo, M. et al. Synaptotagmin 13 is neuroprotective across motor neuron diseases. Acta Neuropathol. 139, 837–853 (2020).

39. Kanda, M. et al. Synaptotagmin XIII expression and peritoneal metastasis in gastric cancer. Br. J. Surg. 105, 1349–1358 (2018).

40. Li, Q., Zhang, S., Hu, M., Xu, M. & Jiang, X. Silencing of synaptotagmin 13 inhibits tumor growth through suppressing proliferation and promoting apoptosis of colorectal cancer cells. Int. J. Mol. Med. 45, 237–244 (2020).

41. Zhang, L. et al. Identification SYT13 as a novel biomarker in lung adenocarcinoma. J. Cell. Biochem. 121, 963–973 (2020).

42. Schonhoff, S. E., Giel-Moloney, M. & Leiter, A. B. Neurogenin 3-expressing progenitor cells in the gastrointestinal tract differentiate into both endocrine and non-endocrine cell types. Dev. Biol. 270, 443–454 (2004).

43. Thorens, B. et al. Ins1 Cre knock-in mice for beta cell-specific gene recombination. Diabetologia 58, 558–565 (2015).

44. Jung, B. et al. Novel small molecules targeting ciliary transport of Smoothened and oncogenic Hedgehog pathway activation. Sci. Rep. 6, 22540 (2016).

45. Rezania, A. et al. Reversal of diabetes with insulin-producing cells derived in vitro from human pluripotent stem cells. Nat Biotechnol 32, 1121–1133 (2014).

46. Gloeckner, C. J., Boldt, K. & Ueffing, M. Strep/FLAG tandem affinity purification (SF-TAP) to study protein interactions. Curr. Protoc. Protein Sci. Chapter 19, Unit19.20 (2009).

47. Olsen, J. V. et al. Parts per million mass accuracy on an orbitrap mass spectrometer via lock mass injection into a C-trap. Mol. Cell. Proteomics 4, 2010–21 (2005).

48. Cox, J. & Mann, M. MaxQuant enables high peptide identification rates, individualized p.p.b.-range mass accuracies and proteome-wide protein quantification. Nat. Biotechnol. 26, 1367–72 (2008).

49. Cox, J. et al. A practical guide to the maxquant computational platform for silac-based quantitative proteomics. Nat. Protoc. 4, 698–705 (2009).

50. Tyanova, S. et al. The Perseus computational platform for comprehensive analysis of (prote)omics data. Nature Methods vol. 13 731–40 (2016).

51. Zhou, Y. et al. Metascape provides a biologist-oriented resource for the analysis of systems-level datasets. Nat. Commun. 10, 1523 (2019).

52. Rouser, G., Fleischer, S. & Yamamoto, A. Two dimensional thin layer chromatographic separation of polar lipids and determination of phospholipids by phosphorus analysis of spots. Lipids 5, 494–6 (1970).

53. Kourtzelis, I. et al. DEL-1 promotes macrophage efferocytosis and clearance of inflammation. Nat. Immunol. 20, 40–49 (2019).

54. Wolf, F. A., Angerer, P. & Theis, F. J. SCANPY: Large-scale single-cell gene expression data analysis. Genome Biol. 19, 15 (2018).

55. Wolf, F. A. et al. PAGA: graph abstraction reconciles clustering with trajectory inference through a topology preserving map of single cells. Genome Biol. 20, 59 (2019).

56. Kuleshov, M. V. et al. Enrichr: a comprehensive gene set enrichment analysis web server 2016 update. Nucleic Acids Res. 44, W90–7 (2016).

57. Blondel, V. D., Guillaume, J. L., Lambiotte, R. & Lefebvre, E. Fast unfolding of communities in large networks. J. Stat. Mech. Theory Exp. 10, P10008 (2008).

